# White matter microstructure changes across the lifespan: a meta-analysis of longitudinal diffusion MRI studies

**DOI:** 10.1101/2025.08.15.669078

**Authors:** Karis Colyer-Patel, Jalmar Teeuw, Vivien Maes, Vera Goossens, Rachel M Brouwer, Neda Jahanshad, Paul M Thompson, Hilleke E Hulshoff Pol

## Abstract

**Background:** White matter in the human brain is known to play a critical role in facilitating communication between different brain regions. White matter microstructure is often quantified using fractional anisotropy (FA) derived from diffusion-weighted MRI and is often considered a key measure of neural efficiency that is positively associated with motor and cognitive functioning. While lifespan trajectories of FA have been well studied in cross-sectional designs, it remains less clear how FA changes longitudinally with age across the lifespan, and whether the rates of change are influenced by genetic variation.

**Methods:** We systematically reviewed the evidence of white matter changes, as measured by fractional anisotropy (FA) with diffusion magnetic resonance imaging longitudinally across the lifespan, and the genetic influences on this change. Searches were conducted in Medline, PsycInfo, and EMBASE up to August 2023 with terms related to DTI/FA and longitudinal/change. Following this, genetic-related search terms were applied to the results, and the search was broadened to include other measures of white matter change. Our systematic search resulted in 29 studies that met our criteria. In addition, 14 studies investigated genetic influences on FA change rates across the lifespan. A meta-regression using a thin-plate spline model was conducted to examine annual whole-brain FA change as a function of age.

**Results:** Across childhood and adolescence, FA increased, and the rate of increase slowed into early adulthood. Between ages 20 and 35, changes in FA were not statistically significant. This was followed by a significant decline in FA between ages 36 and 50. The decreases plateaued between ages 51 and 61 and then continued at a slightly slower rate towards the upper end of the age range assessed (77 years). Average FA change per year relative to baseline assessment reached a maximum of +1.1% during development, and-0.6% per year, during ageing.

Significant heritability was found for *change* in local but not global FA during development. During ageing, common variants in genes that have been related to increased risk for neuropsychiatric disorders (*APOE*, *HTT, MAPT*) were associated in some studies with accelerated local FA decreases over time.

**Conclusions:** In conclusion, there are changes in white matter microstructure within individuals across the lifespan, with increases during childhood, adolescence and early adulthood, followed by a period of relative stability during early to mid-adulthood, and subsequent gradual declines from midlife onwards. Evidence is emerging for genetic influences on white matter changes over time, shaping individual trajectories.

## Introduction

Across the lifespan the brain undergoes significant changes, which coincide with the maturation of cognition and behaviour during development, and with the functional decline found during ageing. The human brain’s white matter microstructure has long been known to play a central role in both development and ageing (Nagy et al., 2004; Lebel et al., 2019; Madden et al., 2012). Moreover, it has been associated with risk and resilience to psychiatric and neurological disorders (Meyer & Lee, 2019; Sorond & Gorelick, 2019). While cross-sectional studies have provided valuable insights into how white matter microstructure changes across the lifespan (Zhu et al., 2025; Villalon et al., 2025), it is still not fully understood how these patterns evolve longitudinally. Longitudinal studies provide unique information on change within individuals that is not available in cross-sectional studies (Hulshoff Pol and Brouwer, 2025). In fact, estimates of change from truly longitudinal assessments may differ from those based on cross-sectional, due to selection and attrition biases (Di Biase et al., 2023). Moreover, it is evident that genetic factors influence white matter microstructure across the lifespan based on studies in twins (Bohlken et al, 2014; Kochunov et al., 2014). So far, it is less clear whether *change* rates of white matter microstructure are also influenced by genes, which would be of interest for clinical trials of drugs to promote healthy ageing (either for therapeutic target development or stratification of data). Therefore, the aims of this systematic review are twofold: one, to investigate how white matter microstructure changes across the lifespan, as measured with changes in fractional anisotropy (FA) using a longitudinal design; and two, to review the evidence of genetic influences on change rates in white matter microstructure across the lifespan.

White matter microstructure is often measured by diffusion magnetic resonance imaging (dMRI). Using dMRI data, microstructural connectivity can be quantified in diffusion tensor imaging (DTI) models (Basser et al., 1994). In DTI, diffusion of water molecules is used to provide indices of (brain) tissue integrity. One of the most common measures of white matter microstructure is FA, which measures the coherence of the orientation of water diffusion, independently of rate (Bennett, 2010). FA is sensitive to myelination, axonal packing, and axon coherence and diameter (Beaulieu, 2002). FA is a robust measure of overall integrity/directionality and is highly sensitive to general connectivity changes (Zhao et al, 2021). FA values can range from 0-1, with higher FA values corresponding to a more consistent diffusion orientation. and often considered to be reflecting higher white matter integrity/directionality. Reductions in FA have been suggested to reflect loss of myelin or axonal degeneration, corresponding to reduced white matter integrity/directionality (Alexander et al., 2011). We note that white matter integrity has frequently been used to refer to FA. To acknowledge that change in FA may reflect multiple changes in white matter, throughout the text we refer to changes in white matter microstructure.

FA changes across the lifespan have been investigated during development (Lebel et al., 2019) and ageing (MacDonald & Pike, 2021), but there are limited studies investigating full lifespan trajectories. Of the few studies that have investigated FA change across the lifespan, most evidence comes from cross-sectional studies (Lebel et al., 2012; Zhu et al., 2025; Villalon et al., 2025). Consistent cross-sectional findings across studies reveal FA values to increase rapidly across the first year of life continuing at a slower pace into early childhood and adolescence (Lebel & Deoni, 2018). During adulthood, FA is more stable, followed by decreases during ageing (Bisdas et al., 2008; Camara et al., 2007; Sullivan & Pfefferbaum, 2006). Longitudinally, in a recent systematic review and meta-analysis, significant within-person decreases in FA were found during adulthood (Mendez Colmenares et al., 2023). We build on this by investigating within-person changes in white matter microstructure across the lifespan (as opposed to focusing on adulthood alone) and by including both healthy individuals and patient cohorts, as well as including genetics of FA change.

The current systematic review and meta-analysis evaluates the evidence up to August 2023 regarding FA changes across the lifespan, by integrating longitudinal findings from individual studies. Only longitudinal studies are included to ensure that individual trajectories of change can be identified. We focus on whole-brain FA changes, but we do include secondary analyses investigating change rates of FA in individual fibres. For a comprehensive overview of genetic influences on change rates of white matter microstructure across the lifespan based on longitudinal studies, twin studies, GWAS, linkage and candidate gene studies are included. By tackling these questions, we aimed to gain a greater understanding of how white matter microstructure changes longitudinally across the lifespan, as well as understanding the role of genetic variations in these changes. Ultimately, these efforts will potentially enable us to understand the biological mechanisms that influence brain development and ageing, as well as those linked to disease.

## Method

### Search strategy

The Preferred Reporting Items for Systematic Reviews and Meta-Analyses (PRISMA) guidelines were followed for the current systematic review (The PRISMA Group, 2020), including for the reporting of study characteristics and findings of included articles. Initial searches were conducted in Medline, PsycInfo, EMBASE, Web of Science Core Collection and Scopus during August 2023 with terms related to structural connectivity/DTI and longitudinal/change (see Appendix for full search strategy and syntax). To review the evidence on influences of genes on white matter microstructure change, we conducted an additional search adding the following search terms: i.e., genes, genetics, genome-wide association study, polygenic, twins and heritability, as well as broadening our search to include other measures of white matter change, for example, local and global efficiency.

### Study selection

One author (KCP) selected, based on all search results, whether the retrieved studies met the inclusion criteria. First, only titles and abstracts were screened and the articles that did not meet the inclusion criteria were excluded. Next, the full texts of the remaining articles were reviewed and the articles that did not meet the inclusion criteria were excluded. Three authors (JT, HHP & VG) conducted a partial screen of the studies and independently reviewed a random sample of 5 studies to confirm that there was agreement between the authors in terms of study selection. The inclusion criteria were: 1) human samples of healthy and/or patient groups (minimum N ≥ 25 per group with longitudinal data/with at least two repeated assessments) in the age range of 0-99 years old; 2) a follow-up duration for repeated MRI scans of at least 6 months; 3) whole brain FA analysis or studies that measured FA in a number of tracts that equated to at least half of the brain’s white matter tracts; 4) primary quantitative data collection (i.e., no case studies or review papers); 5) written in English; 6) studies published in a peer-reviewed journal before August 9, 2023 (**Figure 1**). Only one paper was included if the same cohort was included across more than one study. The study that was included was based on the largest N, largest interval or largest number of assessments. Studies involving a sample group exposed to a drug were excluded, except when a control group was included; in such cases, only the control group data were retained.

**Figure 1.**
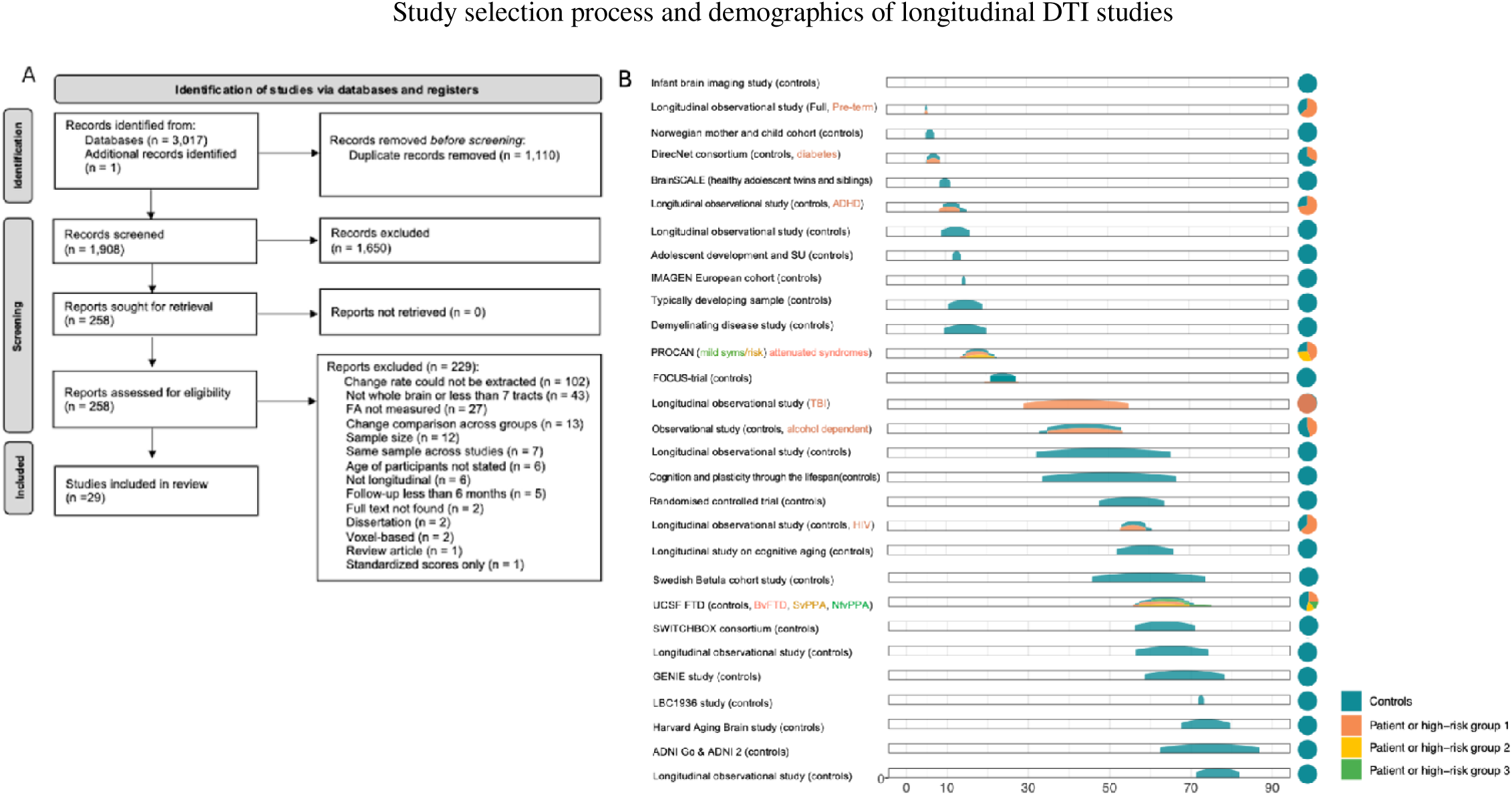
A. PRISMA flow diagram detailing the screening process for the meta-analysis. **B**. Overview of demographics of included studies. Per cohort, an age distribution is displayed based on mean and standard deviation of the age at baseline. On the right, the total number of included subjects is displayed and a pie-chart of the distribution of controls/individuals not belonging to a diagnostic group (blue), and cohorts including one (in orange), two (yellow) or three (green) high risk/diagnostic groups. Abbreviations: ADHD Attention Deficit Hyperactivity Disorder, BP Bipolar disorder, BvFTD Behavioural variant frontotemporal dementia, mild syms Mild Symptoms of attenuated syndromes, NfvPPA Non-fluent variant of primary progressive aphasia, risk Familial risk of attenuated syndromes, SU Substance use, SvPPA Semantic variant of primary progressive aphasia, TBI Traumatic Brain Injury, HIV Human Immunodeficiency Virus.

Our systematic search resulted in 1,908 studies once duplicates were removed. After an initial review of titles and abstracts, 1,650 studies were excluded, primarily due to irrelevant study topics or failure to meet one of the inclusion criteria, such as measuring FA, having a longitudinal design or meeting the specified sample size. A full-text review was then conducted for 258 studies, resulting in the exclusion of 229 studies (see flow diagram in **Figure 1)**. Of note, from the 258 studies that were full text reviewed, 127 studies measured FA across two timepoints. From these studies, 20 studies had quantified whole-brain FA change and so were included. An additional 9 studies quantified FA change in a number of tracts that equated to at least half of the brain’s white matter tracts, leading to the inclusion of 29 studies in total (**Figure 1; Tables 1**). In most studies, DTI acquisition involved a single-shot diffusion-weighted spin echo planar imaging sequence. The scanner type varied across studies, including GE, Philips, or Siemens models, with scanner strength at either 1.5 or 3 tesla (**Supplementary 1.1**).

**Table 1.**
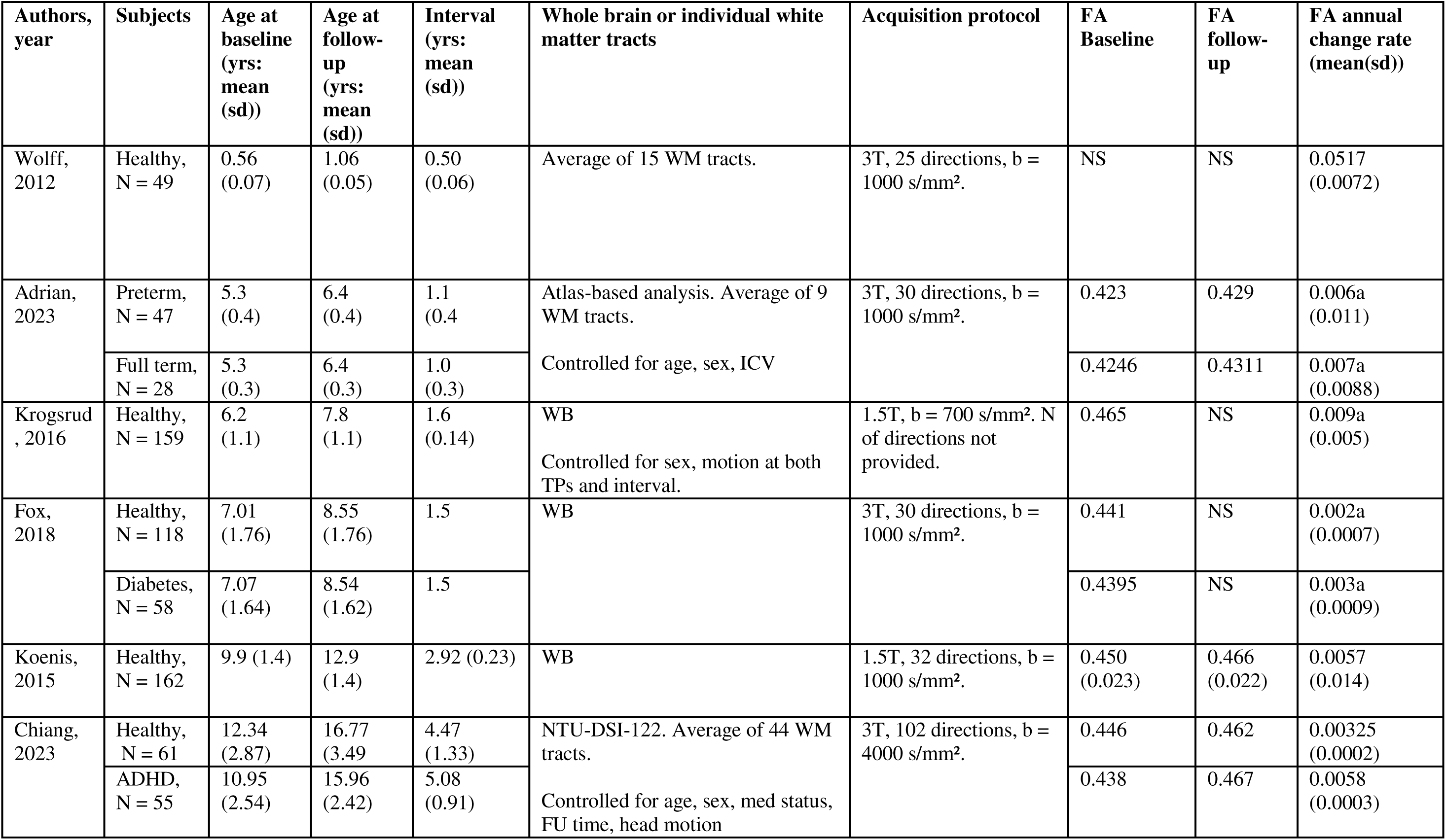
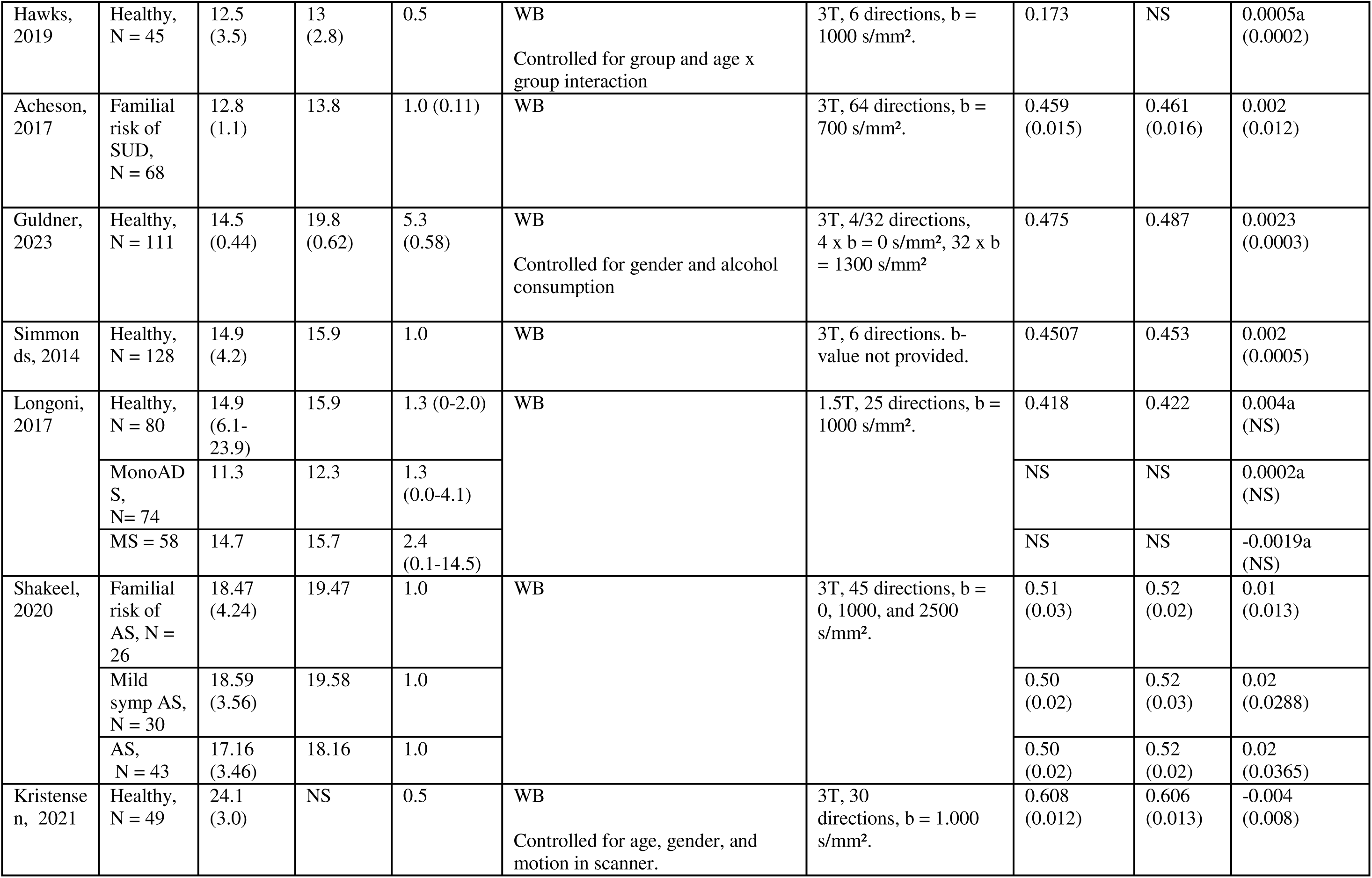
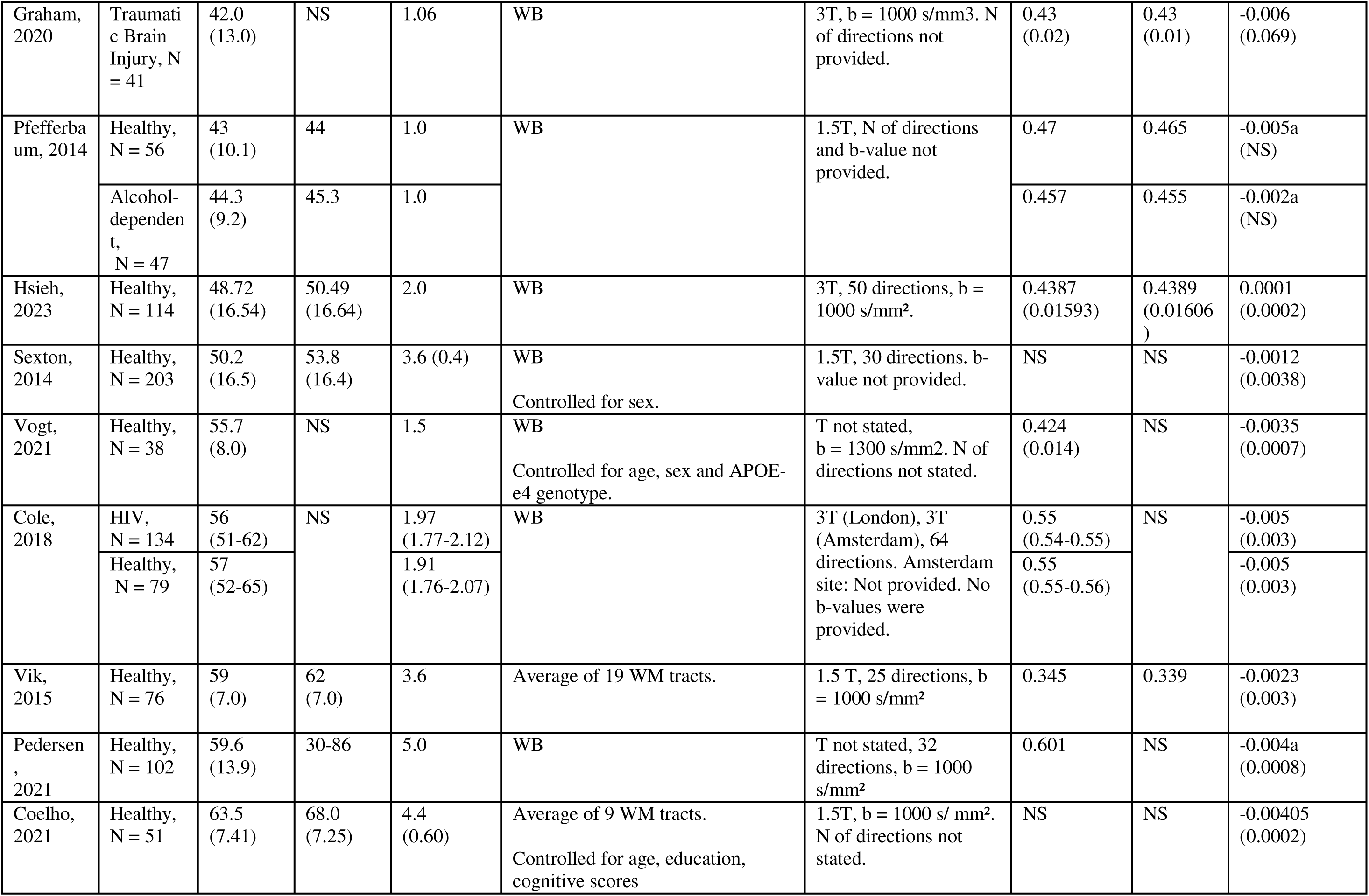
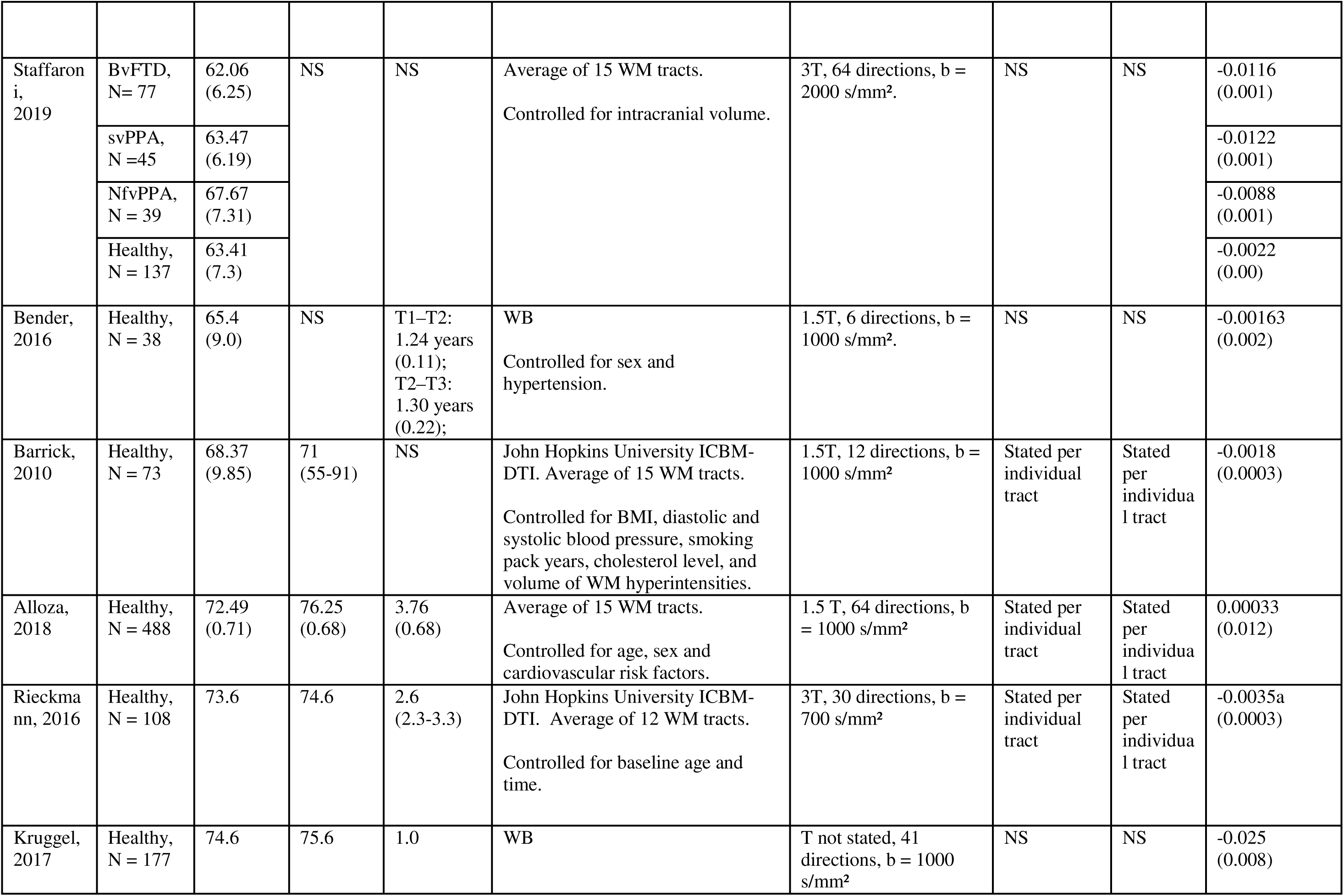
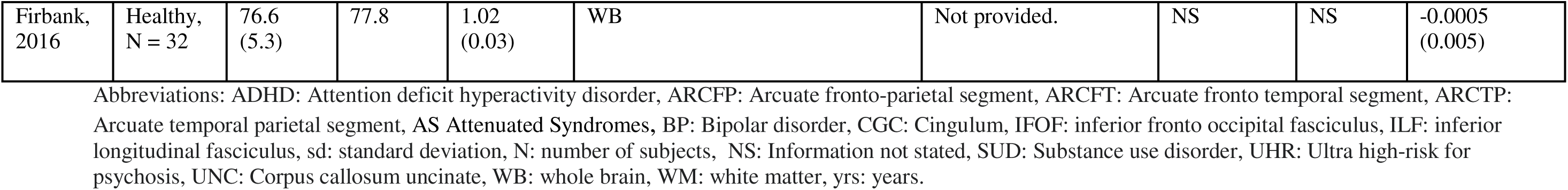
Longitudinal DTI studies measuring FA-change across the lifespan in humans, ordered by age.

Based on the initial DTI study search, twins, GWAS, genes, genetics, polygenic and heritability were added as search terms. In addition, the search was broadened to include other measures of white matter change, for example, local and global efficiency. Fourteen studies were identified as suitable for the systematic review **(Tables 2 and 3).** Please note that only three of these genetic studies were included in the meta-analyses. The remaining studies were excluded due to the unavailability of the data required to extract or estimate FA change rates.

Study selection process and demographics of longitudinal DTI studies

**Table 2.** Longitudinal DTI studies measuring FA change in twins.

| **Authors, year** | **Subjects** | **Age at baseline (yrs: mean (sd))** | **Age at follow-up (yrs: mean (sd))** | **Interval (yrs: mean (sd))** | **Brain region** | **Results** |
| --- | --- | --- | --- | --- | --- | --- |
| Chen, 2009 | N = 60: 30 pairs of same sex twins<br>Neonates | Close after birth | 1 year | 1.0 | Whole brain FA<br>Voxel-based analysis | ↑FA over time.<br>Significant heritability for FA was found in left parietal white matter<br>No significant heritability of change in FA. |
| Brouwer, 2012 | N = 203<br>MZ = 88<br>DZ = 115<br>BRAINSCALE | 9.2 (0.1) mean age for full sample | 12.1 (0.3) | 2.9 (0.2) | 14 ROIs: AF, UF, fornix, ILF, IFOF, GCC, SCC<br>Controlled for multiple comparisons | All tracts ↑FA over time (+1.3% to 6.9%). Cingulum change was most pronounced (6.9%).<br>The heritability of change in mean FA, indicated that the change was not likely influenced by genes. |
| Koenis, 2015 | N=203<br>MZ=88<br>DZ=115<br>BRAINSCALE | 9.2 (0.1) mean age for full sample | 12.1 (0.3) | 2.9 (0.2) | FA and FA-based network changes focussing on global and local efficiency | ↑FA over time for global FA, global efficiency and local efficiency by 3.8%, 3,6% and 3.9%.<br>Heritability was 32% for global efficiency and up to 74% for local efficiency at measurement one, and 48% and up to 75% at follow up.<br>Over time, a stable genetic factor influenced global (genetic correlation ( $r_g$ )=1.00 [0.63–1.00]) and average local ( $r_g=1.00$ [0.53–1.00]) efficiency. |
| Lee, 2019 | N=215<br>74 MZ; 88<br>DZ<br>Neonates | 1 | 2 | 1 | Whole brain FA and<br>in 34 tracts | ↑FA over time.<br>Heritability of FA was $h^2=72\%$ at 1 year of age and $h^2=76\%$ at 2 years of age<br>No significant heritability for change in whole brain FA. Significant heritability of FA change in the right motor segment of the corticothalamic tract and in the right inferior longitudinal fasciculus (ILF) |
| Teeuw, 2022 | N=311<br>214 MZ and<br>DZ twins and<br>97 of their<br>siblings<br>BRAINSscale | $10.0 \pm 1.4$ | 13.0 (1.5)<br>18 (1.4) | 3 and 5 | Global FA and in 20<br>tracts: ATR, CGC, CGH,<br>CST, Fmajor, Fminor,<br>ILF, IFOF, SLFtemp,<br>UNC | ↑FA over time. The maturation of the mean fractional anisotropy of the white matter tracts started to slow down nearing the end of adolescence.<br>Genetics played a considerable role in FA at all ages ($h^2=64\%$; 73%; 51%) and was highly correlated between measurements<br>Heritability of longitudinal change rates in global mean FA during adolescent development (up to $h^2=13\%$ between age 13 and 18 years) was not significant. Locally, FA change was heritable for the cortical spinal tract (38%) and uncinate fasciculus (up to 36%) between 10-18 years, and for the left uncinate fasciculus ($h^2=43\%$) between 13-18 years. |

Abbreviations: ATR: anterior thalamic radiation, CGC: cingulum (cingulate gyrus), CGH: cingulum (hippocampal), CI: confidence interval, CST: corticospinal tract, FA: fractional anisotropy, Fmajor: forceps major, Fminor: forceps minor, ILF: inferior longitudinal fasciculus, IFOF: inferior fronto-occipital fasciculus, SLF: superior longitudinal fasciculus, SLFtemp: superior longitudinal fasciculus (temporal part), UNC: uncinate fasciculus.

**Table 3.** Longitudinal DTI studies on FA change and associations with specific genes and genome-wide findings.

| Authors, year | Subjects | Age at baseline (years: mean (sd) [range]) | Age at follow-up (years: mean (sd)) | Interval (years: mean (sd)) | Genetic variants | Brain region | Results |
| --- | --- | --- | --- | --- | --- | --- | --- |
| Rieckman n, 2016 | N = 108 of which APOE4+ (at least one e4 allele) carriers: 28.7% = 31 | 73.7 [66-87] | NS | 2.6 | APOE4+ compared to APOE4- (no APOE2) genotype | 12 ROIs: SLF; SFOF; IFOF; ACR; SCR; PCR; IC; GCC; BCC; SCC; CGC; CGH<br>Corrected for multiple comparisons | ↓FA over time<br>APOE4+ carriers compared to APOE4- carriers showed no differences in extent of change in FA over time in any of the ROIs, suggesting no genetic influence of the APOE4 allele on change in WM integrity over time |
| Ritchie, 2017 | N = 731<br>APOE4 carriers (at least 1 e4 allele; estimated at approximately 30% of total) versus non-carriers<br>LBC1936 cohort | 69.5 (0.8) | 2nd: 72.5 (0.7)<br>3rd: 76.3 (0.7) | 2nd: 3<br>3rd: 7 | APOE4 genotype | 12 ROIs: GCC, SCC, CGC, AF, UF, IFOF, ATR<br>Corrected for multiple comparisons | At baseline, APOE4 carriers as compared to non-carriers had no differences in FA in any of the ROIs<br>Overall, ↓FA over time<br>APOE4 carriers showed no change in FA over time in any of the ROIs, suggesting no significant genetic influence of the APOE4 allele on change in WM integrity over time |
| Shaffer, 2017 | $N = 261$<br>Pre-Huntington HTT genotype = 191<br>Low CAP = 50<br>Med CAP = 56<br>High CAP = 85<br>Non-carriers = 70 | Low CAP: 36.3 (8.8)<br>Med CAP: 44.0 (10.8)<br>High CAP: 51.2 (11.9)<br>Con = 47.9 (12.1) | Not provided. | Low CAP: 1.4 (1.4)<br>Med CAP: 1.5 (1.5)<br>High CAP: 1.4 (1.4)<br>Con: 1.5 (1.4) | HTT genotype | 9 tracts: premotor area-caudate, premotor area-putamen, right primary motor-caudate, primary motor cortex-putamen, sensory cortex-putamen<br>Corrected for multiple comparisons | At baseline, High CAP versus Low CAP and versus non-carriers: in right primary motor cortex-caudate, and in bilateral premotor area-caudate tracts; ↓FA in left premotor area-putamen tract (versus Con only); Medium CAP versus non-carrier: ↓FA in left premotor area-caudate tract<br>Over time, High CAP versus Low CAP and non-carriers: ↓FA over time in left premotor area-putamen; ↓FA over time in the right premotor area-caudate (versus non-carrier only)<br>Overall, decreases in WM integrity over time are most severe in high CAP groups, closest to transition to HD. |
| Alloza, 2018 | szPRS = 731<br>LBC1936 cohort | 69.5 (0.8) | 2nd: 72.5 (0.7)<br>3rd: 76.3 (0.7) | 2nd: 3<br>3th: 7 | szPRS | 12 ROIs: GCC; SCC; and bilateral C; ATR; AF; UF; ILF<br>FA and graph metrics<br>Corrected for multiple comparisons | No significant associations with szPRS and FA and graph metrics at baseline<br>↓FA over time, irrespective of szPRS<br>szPRS did not show significant associations with FA change or graph metrics over time. This suggests that szPRS may not directly influence change in FA over time. |
| Swanson, 2018 | $N = 100$<br>FXS = 27 | FXS: 6.5 (0.8) months | 2nd: FXS: 12.6 (0.8) months | 2<sup>nd</sup>: 6 months<br>3<sup>rd</sup>: 18 | FXS | 19 ROIs: the bilateral ALIC, PLIC, ATR, ILF, UF, SCP, MCP, and 6 segments of the CC | FXS versus non-carriers: ↓FA in bilateral ALIC, the ILF, the SCP, UF and 4 segments |
|  | Non-carriers = 73 | Con: 6.7 (0.7) months | Con: 12.6 (0.6) months<br>3rd: FXS: 24.7 (1.0)<br>Con: 24.8 (1.3) months | months |  | Corrected for multiple comparisons | of the CC.<br>FXS and non-carriers both have ↓FA over time.<br>No differences in FXS versus non-carriers in FA-change over time. This suggests no genetic influence of FXS on change in FA over time. |
| Panman, 2019 | *N* = 113<br>Presymptomatic C9orf72 repeat expansion = 12<br>Presymptomatic MAPT mutation = 15<br>GRN mutation = 33<br>Non-Carriers = 53 | C9orf72 : 49.7 (12.4)<br>MAPT: 41.8 (9.5)<br>GRN: 52.1 (7.5)<br>Non-carriers: 50.7 (10.7) | NS | 2 | C9orf72 repeat expansion<br>MAPT mutation<br>GRN mutation | Whole brain FA analysis<br>Corrected for multiple comparisons | At baseline, lower FA in C9orf72 repeat expansion carriers in frontotemporal tracts versus non-carriers, predominantly located in the bilateral corticospinal tract and anterior thalamic radiation, the right inferior fronto-occipital fasciculus, and superior longitudinal fasciculus. At baseline, no differences in MAPT and GRN carriers compared to Non Carriers.<br>C9orf72 and GRN carriers showed no significant changes in FA over time versus non-carriers.<br>MAPT carriers ↓FA over time in the left UF, the left ATR, and the left IFOF, versus non-carriers.<br>Overall, this suggests that MAPT mutation influences ↓FA over time, while C9orf72 repeat expansion and GRN mutation do not. |
| Williams, 2020 | $N = 406$<br>presymptomatic APOE4 carriers = 96<br>Non Carriers = 310 | 71.3 (9.9) [50-95] mean age for full sample | 75 | 3.6 (1.7) [0-8.3] | APOE4 genotype | 16 ROIs: ACR, ALIC, PLIC, BCC, GCC, CGC, CFX, BFX, FX/ST, IFOF, SFOF, PCR, SCR, SLF, and SS | APOE ε4 carriers versus Non Carriers did not show any significant differences in FA at baseline<br>Overall, ↓FA over time, except for GCC and posterior limb internal capsule<br>APOE4 carriers versus non-carriers: ↓FA over time in the GCC and SCC. This suggests genetic influence of APOE4 on FA-change over time |
| Bagautdinova, 2020 | $N = 201$<br>22q11.2DS = 101<br>Non-carriers = 100 | 22q11.2DS: 15.9 (6.0)<br>Con: 16.6 (6.8) | 2nd 22q11.2D S: 20 Con: 20<br>3rd 22q11.2D S: 23 Con: 24 | 1st: 3.7 (0.89)<br>2nd: 3.3 (0.8) | 22q11.2DS | 18 ROIs: ATR, CST, ACB, CGC, ILF, SLF-parietal bundle, SLF-temporal bundle, UF bilaterally, FMj and FMi (CC)<br>Corrected for multiple comparisons | 22q11.2DS show consistently ↑FA vs non-carriers in bilateral ATR, CGC, SLF-parietal bundle, FMj and FMi (CC)<br>Both 22q11.2DS and non-carriers have ↑FA over time<br>No significant differences in change in FA over time in 22q11.2DS vs non-carriers. This suggests no genetic influence of 22q11.2DS on FA-change over time |
| Elsheikh, 2020 | $N = 137$<br>AHD = 31<br>MCI = 57<br>Non-carriers = 49 | AHD: 76.5 (7.4)<br>MCI: 75.3 (5.9) | 2nd AHD: 77<br>MCI: 76<br>HC: 78 | 1 | AHD<br>MCI<br>GWAS | Whole brain global connectivity, i.e., Louvain modularity, transitivity, global efficiency and characteristic path length | AHD had changes in global connectivity over time in Louvain modularity, global efficiency and path length. MCI and non-carriers had no change in connectivity over time.<br>Louvain modularity was associated with CDH18 and almost reached genome wide significance |
|  | ADNI cohort | HC:<br>77.0<br>(5.1) |  |  |  | Corrected for multiple comparisons GWAS (threshold for 5% set at 5e-6) | Transitivity, global efficiency and characteristic path length had associations with several other genes, including with OR5L1, OR5D13, OR5D14, IGF1 and more, at the adapted threshold of 5e-6 |

### Data extraction for studies reporting on FA change

For each study, the mean age of the sample was taken as the mean age at timepoint 1 (baseline assessment). Next, mean whole brain FA annual change was extracted from the studies. In some cases, the FA value from timepoint 1 and timepoint 2 (follow-up assessment) were available in which case FA at timepoint 1 was subtracted from FA at timepoint 2. The change was divided by the average follow-up duration in years, to create an annual change rate. In other cases, the annual change rate was explicitly stated and included. For both, measurements were annotated as the Delta (Δ). Studies that investigated whole brain FA changes, but had not provided the exact FA values, were also included. For these additional studies, FA values or FA change rates were estimated based on figures published in these studies. The decision to include studies with missing statistical information (and so this information was estimated) was made based on evidence that doing so can increase the robustness of meta-analyses by allowing a larger number of studies to be included, despite the uncertainty associated with approximated values (Idris & Robertson, 2009; Nieminen, 2022).

In addition, we also extracted any measures of sample variance of the change rates (in SD). For five studies, SD was provided for annual FA change rates. For one study, standard error (SE) was provided, and converted into SD. The SD for the remaining studies was approximated by deriving Z-scores from reported or estimated p-values, which were then used to back-calculate SDs from the reported mean FA changes; see **Supplementary Information 1.2** for full details. Following this, the sampling variance (*vi*) was calculated using the *escalc* function from the *metafor* package (Viechtbauer, 2010) in RStudio version 1.3.1073 (RStudio Team, 2020; http://www.rstudio.com), based on the mean annual FA change, SD, and sample size.

For the studies where whole-brain FA could be extracted or estimated, if the study included more than one group, mean estimates and SD were pooled across these groups to obtain a single estimate per study. This was also repeated for studies that measured FA in a number of white matter tracts covering at least half of the brain’s major pathways. For these studies, we calculated a pooled estimate of annual FA change across all reported tracts. A weighted pooled standard deviation was also computed, and both the estimate and SD were subsequently pooled across groups within each study. Subsequently, the studies were included in the meta-regression. Recent studies on the large population cohorts (ABCD, UK Biobank) were made available after the cut-off date of the search and so were included as validation of the age meta-regression models (Beck et al., 2023; Korbmacher et al., 2024).

### Meta-analysis of annual whole-brain FA change

To examine the relationship between age and annual change in FA, a mixed-effects meta-regression model with restricted maximum likelihood (REML) estimation was fitted using the *rma* function from the *metafor* package (Viechtbauer, 2010). The model included a thin-plate spline term to flexibly capture nonlinear associations between study-level mean age (moderator) and FA change per year. Study-level sampling variances were incorporated as weights, and random effects accounted for between-study heterogeneity. Models were fitted with a spline basis with three to five knots. The final model with the optimal number of knots was selected based on the lowest Akaike Information Criterion (AIC) to balance model complexity and overall fit to the data. Predicted FA change values across the lifespan (ages 6–77 years) were generated using a prediction matrix and used for visualisation. Additionally, the annual percentage change in FA was estimated using the formula: *Percentage chang*e = (Annual Change / Initial FA) × 100. This analysis was repeated for healthy controls only, and for studies where measurements for individual tracts could be extracted.

To estimate actual FA values at specific ages, predicted annual change rates from the spline-based meta-regression were integrated across age using a Riemann sum approximation. This approach reconstructs the FA trajectory by summing predicted changes over time in small steps (0.01 per year), effectively integrating the derivative function estimated by the model. The resulting curve was anchored at age 43 using a reference FA value from Pfefferbaum et al. (2014), selected because the sample’s mean age closely matched the average age across studies included in the meta-analysis. Additionally, the same method was repeated for the corpus callosum. The resulting curve during development was anchored at age 11.68 using a reference FA value from Chiang et al., (2023), and during ageing was anchored at age 72.49 using a reference FA value from Alloza et al., (2018).

### Sensitivity analysis

Several sensitivity analyses were conducted to assess the robustness of the meta-regression findings. Firstly, to evaluate the influence of potential outliers, we successively excluded three studies that appeared report extreme values relative to the rest of the studies and single-handedly exerted heavy influence on the curve of the age-dependent trajectory (Kruggel et al., 2017; Shakeel et al., 2020; Wolff et al., 2012) **(Supplementary 1.3; Figure S1)**.

Secondly, to evaluate the impact of including studies with approximated parameters, a second sensitivity analysis was performed. Studies were classified into four categories based on the availability and completeness of their reported statistical information. Category 1 [ideal] included studies that reported both a point estimate and either an SD or a p-value. Category 2 comprised studies where the point estimate was approximated, but the p-value was reported and used to estimate the SD. In Category 3, studies provided a point estimate, but the SD had to be estimated using a conservative p-value. Finally, Category 4 [less ideal] included studies for which both the point estimate and the SD or p-value had to be approximated. Each category was added in succession to assess whether the trajectory of FA change varied.

Third, we examined whether including studies that measured individual tracts covering at least half of the brain (Category 5), allowing a proxy for whole-brain FA to be calculated, influenced the trajectory, compared to those that measured whole-brain FA directly.

Finally, we conducted a sensitivity analysis excluding studies that included patient groups to assess whether patients with a neurological and (neuro)psychiatric condition affected the curvature of the lifespan trajectory.

## Results

The model with three knots provided the best fit, as indicated by the lowest AIC compared to the models with four and five knots (**Supplementary Table S1**). The meta-analysis revealed significant FA changes with age across longitudinal DTI studies (**Figure 2; Supplementary Information 1.4; Supplementary Table S2**). This was reflected in the estimated trajectory of whole brain FA **(Figure 2b).** Across childhood and adolescence, FA increased, with the rate of increase slowing into early adulthood. Between approximately ages 20 and 35, changes in FA were not statistically significant. This was followed by a decline in FA between ages 36 and 50. The decreases plateaued between ages 51 and 61, and then continued at a slightly slower rate towards the upper end of the age range (77 years). Average FA change per year relative to baseline assessment during development was up a maximum of +1.1%, and a minimum of-0.6% during ageing.

**Figure 2.**
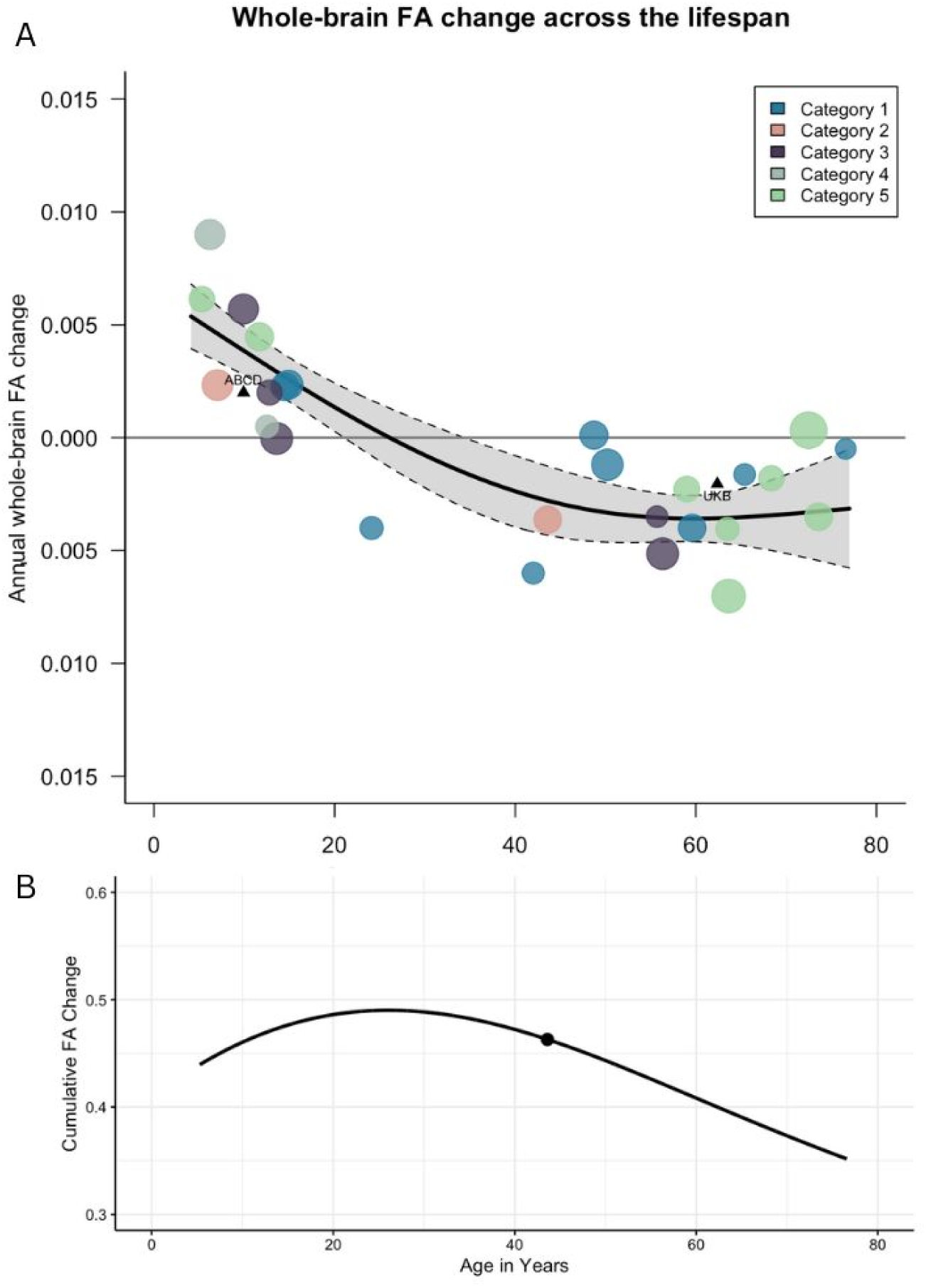
**Meta-analysis of whole-brain FA change across the lifespan**. Annual FA change per year represents the FA at timepoint 2 minus FA at timepoint 1, divided by the number of years between timepoints. Each circle represents a cohort, with circle size proportional to cohort size (ranging from N = 32 to N = 213). Data were pooled across groups within each study. Studies were classified into four categories: Category 1 reported both point estimates and SD or p-values (blue circles); Category 2 required approximation of the point estimate but reported p-values to estimate SD (green circles); Category 3 reported point estimates but used a conservative p-value to estimate SD (purple circles); Category 4 required approximation of both the point estimate and SD or p-values (grey circles); Category 5 included studies that measured FA change across individual tracts, from which a pooled estimate of annual FA change was calculated. **A.** A thin-plate spline model was used to fit the data, shown with 95% confidence intervals. **B.** The estimated trajectory of whole-brain FA values across the lifespan was derived by integrating the predicted annual FA change rates from the spline model using Riemann sum approximation. The trajectory was anchored at age 43 using a reference FA value from Pfefferbaum et al. (2014), which is represented by a black dot to reflect the average age of samples included in the meta-analysis.

Sensitivity analysis indicated that including different categories of studies, based on whether data could be directly extracted or estimated, did not affect the overall results. Therefore, all four categories were retained in the main analysis (**Figure 2; Supplementary Information 1.5; Figure S2; Supplementary Table S1**). We also found that including studies that measured individual tracts (from which a proxy for whole-brain could be calculated) did not affect the overall trajectory, and these were therefore retained as well (**Figure 2; Supplementary Information 1.6; Figure S3; Supplementary Table S1**).

Finally, excluding patient groups did not have a large impact on the curvature. However, confidence intervals were smaller around the transition from increasing to non-significant change, as well as towards the upper end of the age range (see **Supplementary Information 1.7; Figure S4**).

We also investigated annual FA changes across the lifespan on an individual tract level. Across studies, measurements were mostly available for the frontal and temporal tracts, as well as the inferior frontal-occipital fasciculus, inferior and superior longitudinal fasciculus, cingulum, uncinate fasciculus and internal and external capsule. Additionally, for the *corona radiata*, thalamic radiation, corticospinal, fornix and corpus callosum (**Supplementary Table S3)**.

However, we faced difficulty when fitting the meta-analytic models due to the limited data being available across adulthood (ages 20-55) (**Figure S5**); except for the corpus callosum, where separate models could be fitted to the two age intervals in development (5-12) and advanced adulthood (63-74) with sufficient data available (**Figure 3**). Similarly to whole-brain FA, increases in FA for corpus callosum were found up until early adolescence. Due to the relative lack of data between ages of 13-62 whole lifespan changes in FA cannot be estimated. From the ages of 63-70 decreases in FA were found, followed with reductions in the rate of decrease up until the upper age limit of 74.

**Figure 3.**
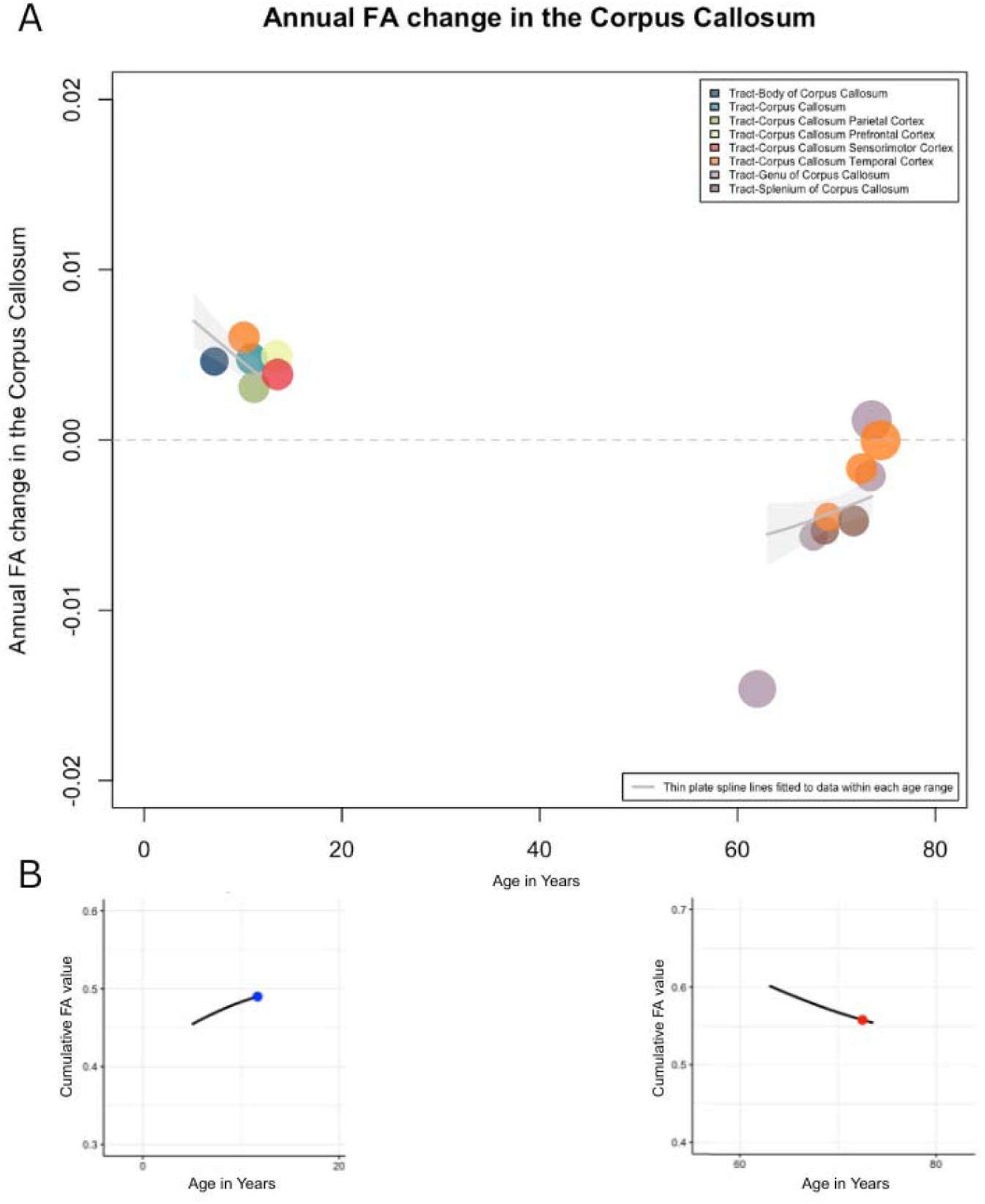
Meta-analysis of FA change rate in the corpus callosum. Annual FA change per year represents the FA at timepoint 2 minus FA at timepoint 1, divided by the number of years between timepoints. Each circle represents a cohort; circle size reflects cohort size (ranging from N = 73 to 427 individuals). Colours indicate different subregions of the corpus callosum. Data were pooled across groups within each study that measured the same corpus callosum region. Notably, the included studies include both healthy and clinical populations. Thin plate spline lines were fitted to the data within two distinct age ranges (5-12 years and 63-74 years) and are shown with their corresponding 95% confidence intervals. **B.** The estimated trajectory of whole-brain FA values across the lifespan was derived by integrating the predicted annual FA change rates from the spline model using Riemann sum approximation. The trajectory during development was anchored at age 11.68 using a reference FA value from Chiang et al., (2023), which is represented by a blue dot. The trajectory during ageing was anchored at age 72.49 using a reference FA value from Alloza et al., (2018), which is represented by a red dot.

### Genetic influences on white matter microstructure changes across the lifespan

In twins, longitudinal studies of heritability (h^2^) for FA change are based on two cohorts so far, namely one in neonates and one in emerging adolescence (**Table 2**) (Lee et al., 2019; Teeuw et al., 2022). The heritability for change in whole-brain FA over time and changes in global and local efficiency over time did not reach significance in these two cohorts. However, local/tract-specific longitudinal changes over time in FA did reveal significant influence of genetic factors in the right motor segment of the corticothalamic tract and in the right inferior longitudinal fasciculus (ILF) between 0-1 year of age (Lee et al., 2019), and in cortical spinal tract (h^2^=38%) and uncinate fasciculus (h^2^ up to 36%) between 10-18 years of age, and in the left uncinate fasciculus (h^2^=43%) between 13-18 years (Teeuw et al, 2022).

Nine longitudinal studies on genetic variants influencing FA change were retrieved. Eight of these have focussed on specific variants related to disease, including Alzheimer’s disease (*APOE*), Huntington’s disease (*HTT*), Fragile X Syndrome, 22q11.2 Deletion Syndrome, ALS (*C9orf72*, *MAPT*, *GRN*) and schizophrenia (polygenic risk) (**Table 3**). The 9th study reported on a genome-wide association study in a cohort enriched for Alzheimer’s disease and mild cognitive impairment (**Table 3**). Influences of genes on change in FA over time were found in four of those nine studies: for presymptomatic *APOE* ε4 carriers as compared to non-carriers decreased FA over time in the genu and had more pronounced decrease in FA over time in the splenium of the corpus callosum; for *MAPT* mutation carriers compared to non-carriers there was decreased FA over time in the left uncinate fasciculus, anterior thalamic radiation, and inferior fronto-temporal fasciculus; and for pre-Huntington mutant-*HTT* genotype carriers, most pronounced in those closest in time to disease transition, compared to non-carriers, showed decreased FA over time in the left premotor area-putamen and right premotor area-caudate tracts. Given that brain related traits – like cross-sectional FA itself (Zhao et al., 2020) – and their longitudinal change rates are likely to be polygenic in nature, genome wide studies are needed to detect the biological pathways associated with these changes in healthy development and ageing. In the first genome-wide association study on *changes* in FA-based structural brain connectivity over time weighted global efficiency was significantly influenced by the *IGF1* gene on chromosome 12, and characteristic path length was significantly influenced by the *ZDHHC12* and *ENDOG* genes on chromosome 9. In the five other studies no significant influences were found.

## Discussion

This systematic review and meta-analysis evaluated the evidence from longitudinal studies to date of annual whole-brain FA changes across the lifespan in healthy individuals, high-risk and clinical groups. White matter microstructure changed significantly within individuals across the lifespan, with increases in FA during development, stability during young and middle adulthood, and decreases in FA during ageing. More specifically, across childhood and adolescence, FA increased, with the rate of increase slowing into early adulthood. Between approximately ages 20 and 35, changes in FA were not statistically significant. This was followed by a decline in FA between ages 36 and 50. The decreases plateaued between ages 51 and 61, and then appeared to continue at a slightly slower rate towards the upper end of the age range (77 years). Excluding patient groups did not have a large impact on the trajectory of FA change, although this is likely due to the patient groups making up ∼10% of the total sample. Longitudinal change in FA of the corpus callosum followed a similar trajectory to whole-brain FA during both development and ageing. However, in later life, the rate of decline in the corpus callosum appeared to attenuate more sharply than in the whole brain. Currently available longitudinal FA data did not allow for quantifying trajectories to firmly establish heterochronicity in peak FA between separate fibers. Overall, significant heritability for change in local FA is found during early development and during adolescence, but not for global change in FA. Specific genetic variants related to increased risk for neuropsychiatric disorders (*APOE*, *HTT*, *MAPT*) associated with FA change over time, also in the pre-clinical phase. Thus, evidence is emerging that genetic factors are implicated in the human brain’s white matter microstructure changes throughout life.

Longitudinal changes in FA have been associated with changes in motor and cognitive functioning in several studies. For instance, in adults, training of a complex visuomotor skill (juggling) increased FA over time (Scholz et al., 2009). Moreover, increased physical training of an overlearned sensory-motor skill (biking) revealed increases in FA along several tracts, including the cortico-spinal tract connecting the arms and legs with the sensory-motor cortex involved in sensory motor activation (Svatkova et al., 2015). Others revealed similar changes over time based on physical activity changes in children (Chaddock-Heymann et al., 2018).

Moreover, transient changes in FA have been reported during activations in the scanner, including during visual processing and tactile stimulations (Mandl et al., 2013), as well as during loss of consciousness under general anesthesia when compared to before and after anesthesia (Tang et al., 2021). This shows how external or environmental factors might influence change in FA. Also, associations with cognitive functioning have been reported: increases in FA during development have been associated with increases in cognitive functioning (Koenis et al., 2015). In addition, in schizophrenia patients, cognitive dysfunction has been associated with abnormal organization of frontotemporal white matter pathways (Falkai et al., 2023). Finally, decreases in FA during ageing may reflect a lower efficient information transfer and have been associated with a decline in cognitive performance in the ageing brain (Coelho et al., 2021; Damoiseaux, 2017). Thus, FA changes over time coincide with sensory-motor and cognitive functioning during development and ageing and therefore hold relevant functional properties. Indeed, if continued activation of such tissue would result in a strengthening of connections, increase in FA could possibly reflect its use and learning in the longer term.

One of our initial aims was also to investigate FA changes rates on an individual tract level. However, due to the lack of data points covering the lifespan, we faced difficulty when fitting the meta-analytic models for FA change per tract (**Supplementary Figure S5**). An exception was the corpus callosum, for which we were able to model both developmental and ageing-related changes. The trajectory during development and ageing closely mirrored that of whole-brain FA, with the exception of a small difference in later life. The similarity observed during development and the overall decline during ageing were in line with expectations, given that the corpus callosum is the largest white matter tract (Travers et al., 2015). The more pronounced slowing of decline in the corpus callosum may reflect regional differences in vulnerability, as distinct parts of the corpus callosum degrade at different times (Bennett et al, 2010). Alternatively, it may indicate that the corpus callosum undergoes more pronounced age-related decline earlier in life than other tracts. Due to the limited number of available data points, it remains difficult to determine whether our findings provide evidence for differential maturation and degradation of the corpus callosum and its subdivisions. Earlier findings suggest a posterior-to-anterior pattern of development with earlier plateaus found in the splenium compared to the genu of the corpus callosum (Lebel & Deoni, 2018). This would also fit with the general theory that brain regions involved in primary sensorimotor function (e.g., the splenium) tend to mature before higher order association systems (Thompson et al., Nature 2000). Similarly, during ageing the subregions of the corpus callosum are differentially sensitive to degeneration, following an anteroposterior gradient, which is also found across the whole brain (Bennett et al., 2010). A comparison with FA change in other individual fibers did show some suggestive evidence for a more prominent increase in FA during early development for the uncinate and inferior longitudinal fasciculus, and during later development for the cingulum and superior longitudinal fasciculus, while thalamic radiation kept increasing steadily (**Supplementary Figure S5**). During ageing, the most prominent decreases in FA were found in the uncinate and cingulum. These observations could be in line with differential peak FA for separate fibers in the brain. To pinpoint differences in change rates on an individual tract level, and further on a subregion level more longitudinal data is required. Currently, FA tract based longitudinal studies in the age period between 15-60 are limited, as it is often assumed that minimal changes occur during this time. However, individual-level changes may still be present, but these could be obscured by group-level patterns (Hulshoff Pol and Brouwer, 2025) and may be linked to functional outcomes as well as various disorders.

Based on the few studies published using a longitudinal twin study design, significant heritability was reported for change in FA during development in local tracts but not for change in global FA (**Table 2**). This finding could imply that different genes impact change in FA in separate fibers locally in the brain, rather than that there is overlap in genetic variants implicated in multiple fibers throughout the brain resulting in impacting change in FA overall. However, it is also possible that heterochronicity of peak FA differed between fibers, resulting in the sum of FA change throughout the brain being diluted. Indeed, we find that the lifespan trajectory for global mean FA change was flatter (max: +1.1%; min:-0.6%) (Figure 1) than the peaks for development and aging found for mean FA change measured in separate fibers (max: +2.0% for inferior longitudinal fasciculus; min:-2.0% for uncinate fasciculus) (**Supplementary Figure S5**). Heterochronicity of peak FA could also be present within parts of a fiber, such as was recently shown in a GWAS of area and thickness of the corpus callosum, where genetic variants were found for corpus callosum parcellations in a rostral-caudal gradient (Bhatt et al, 2025). We also find that heritability was significant in the right motor segment of the corticothalamic tract and right inferior longitudinal fasciculus during early development (Lee et al, 2019), and during adolescence in the left uncinate fasciculus, among other fibers (cortical spinal tract and uncinate fasciculus) (Teeuw et al, 2022). This finding may be in line with evidence suggesting the left hemisphere develops later than the right hemisphere functionally (Chiron et al, 1997) and in part structurally (Williams et al, 2023). The few longitudinal dMRI studies in individuals at high risk of psychiatric disorders, have focussed on first-degree relatives other than twins, namely siblings or children of patients, where familial risk but no heritability can be established. In individuals at high risk for a psychotic disorder significant changes over time suggest an influence of familial risk (Domen et al., 2017, but see Caspi et al., 2022). In individuals at high risk for bipolar disorder as compared to control individuals, differential connectivity changes over time in a FA-based structural network was revealed on top of considerable development-related changes in all participants (Roberts et al., 2022, but see Ganzola et al., 2017).

Based on cross-sectional twin studies, global estimates of FA heritability average around h^2^∼50% and vary locally between h^2^=20% to 80% (Blokland et al., 2012; Jahanshad et al., 2013; Richmond et al., 2016; Videtta et al., 2024; Kochunov et al., 2014). This can already be measured very early in life with an h^2^=78% reported for FA at age 1 years (Lee et al., 2019) and has been found to remain rather stable during early infant development (Lee et al., 2019). Also, the genetic factors underlying cross-sectional FA overlapped with genetic risk for schizophrenia (Bohlken et al., 2016) and lower cross-sectional FA was found with increased genetic risk for Alzheimer’s disease (Harrison et al, 2020). A genomewide association study (GWAS) on white matter microstructure in 43,802 individuals identified 109 associated loci, 30 of which were detected by tract-specific functional principal components analysis, and common variants associated with white matter microstructure that altered the function of regulatory elements in glial cells, particularly oligodendrocytes (Bhatt et al., 2024). Thus, there is ample evidence for considerable influences of genetics on cross-sectional FA over the lifespan. The heritability of white matter microstructure *change* is much lower than heritability estimates for cross-sectional white matter microstructure per se - although locally, heritability of change in white matter microstructure reached up to h2=43% (Teeuw, 2022). This difference may be expected based on the difference found in heritability for white matter volume change (∼40% in Brouwer et al, 2017) compared to that of white matter volume (∼90% e.g., Baare et al, 2001). Also, the fewer hits found in the longitudinal GWAS on brain volumes (Brouwer et al, 2022) compared to cross sectional GWAS of the same structures (e.g., Hibar et al, 2015) suggests as much, although with longitudinal data being more sparse than cross-sectional data, the number of inclusions may also play a role.

Interestingly, longitudinal dMRI studies on the influence of genetic variants are starting to emerge, and some of these suggest that specific genetic variants related to increased risk for neurological disorders (*HTT*t, *MAPT*, *APOE* allele ε4) are associated with FA change over time also in the pre-clinical phase, albeit less so for psychiatric disorders (**Table 3**). Overall, decreases in FA over time were most severe in high CAP groups (CAG Age Product CAG-repeat length as a proxy for clinical diagnosis probability), closest to transition to Huntington’s Disease (Shaffer et al., 2017). In another study *MAPT* mutation carriers had more pronounced decreases in FA over time in the left Uncinate Fasciculus, the left anterior thalamic radiation (ATR), and the left inferior fronto-occipital fasciculus (IFOF), versus non-carriers. Overall, this suggests that in *MAPT* mutation carriers, a gradual progression of neurodegeneration in white matter microstructure over time takes place (Panman et al, 2019). Moreover, in the first genome-wide association study on structural connectivity change over time, genetic variants implicated in those changes emerged in the Alzheimer’s Disease Neuroimaging Initiative (ADNI) cohort including patients with Alzheimer’s disease, mild cognitive impairment and control individuals (Elsheikh et al., 2020). Thus, while genetic influences on individual differences in cross-sectional white matter microstructure are considerable and likely far exceed those related to FA change over time, some evidence is starting to emerge for genetic variants implicated in *changes* in white matter microstructure over time.

As to the mechanisms explaining the changes in white matter microstructure during development and ageing measured by fractional anisotropy in longitudinal study designs, we can only speculate. Longitudinal changes in FA could reflect higher/lower diffusion-driven displacements of water molecules in one direction (such as along the direction of the white matter fibers) and/or lower/higher diffusion-driven displacements of water molecules in the other directions (such as perpendicular to the white matter fibers) as inferred from the definition of FA (Le Bihan & Iim, 2015). Following spinal cord injury in monkeys, a significant reduction of FA was found above and below injury segments, supporting FA as a most sensitive measure of structural disruptions of white matter microstructure (Misha et al, 2020). FA is now considered to be a sensitive marker to myelin (Lazari and Lipp, 2020), although the degree of myelination does not determine tissue anisotropy since it has also been demonstrated in non-myelinated fibres, such as in the cerebral cortex (Reveley et al., 2022). However, overall, the exact mechanisms as to what the *change* in white matter microstructure represents remains largely unresolved.

In conclusion, white matter microstructure, as measured with FA, was found to change within individuals across the lifespan, with increases during development and decreases during ageing. Limited evidence is reported for heritability of change in FA, as a representation for change in white matter integrity/directionality, during early development and during adolescence.

However, it is important to note that FA is an inherently noisy measure, with only modest test-retest reliability (Chen et al., 2020; Jansen et al., 2007; Buimer et al., 2020). Moreover, specific image processing pipelines have been shown to yield more reproducible results than others (Danielian et al., 2010). As such, large sample sizes, substantial changes over time, or longer follow-up intervals are likely necessary to reliably detect individual differences in change, including genetic effects on the rate of FA change. Nevertheless, there is evidence suggesting specific genetic variants related to increased risk for neurological and psychiatric disorders are associated with FA change over time, also in the pre-clinical phase. To fully elucidate normative development and ageing processes, as well as trajectories associated with various disorders, it is important to measure across a broader range of ages and more diverse clinical groups. Including a higher N will help detect relevant signals (Marek et al., 2022; Spisak et al., 2023), and reporting more detailed statistics of all FA outcomes will ease comparison between studies.

Furthermore, more detailed, possibly higher resolution measurements and other measures such as mean diffusivity and local efficiency should be included (Figley et al., 2022; Sudre et al., 2021). Applying modelling techniques like restriction spectrum imaging to promising higher order acquisitions, such as multi-shell diffusion-weighted imaging (Palmer et al, 2022) and combining these with detailed anatomical atlases (Amunts et al., 2020), will further our understanding.

Including age-dependent associations with genetic variants, as shown earlier for white matter volume change (Brouwer et al, 2022), will help identify finer-grained changes in white matter microstructure with age. Together, these approaches will elucidate which mechanisms are responsible for changes in white matter microstructure and how they affect motor and cognitive functioning during development and ageing.

## Appendix

**Table 4.**
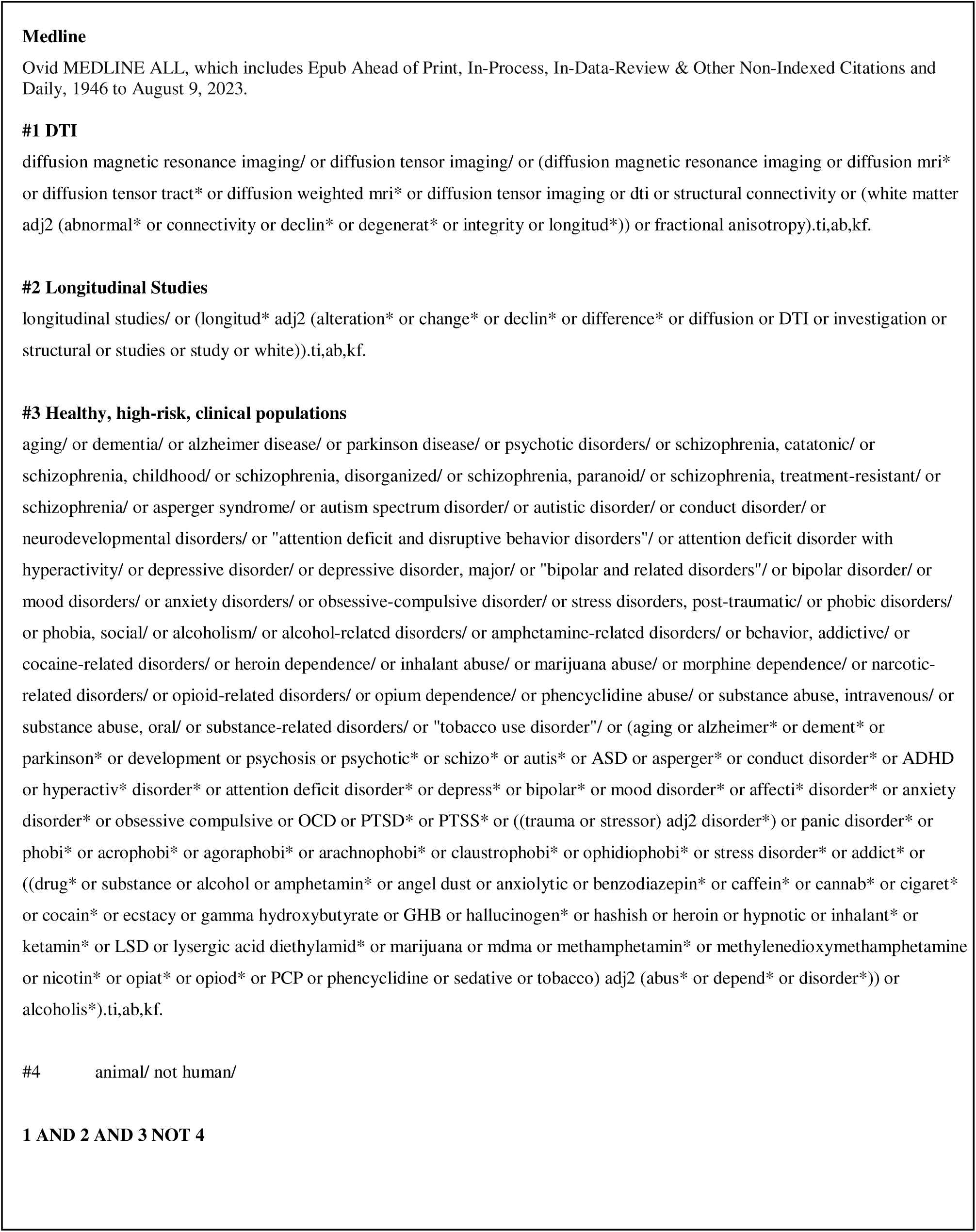
Search syntax from Medline database, last accessed on August 9, 2023.

**Table 5.**
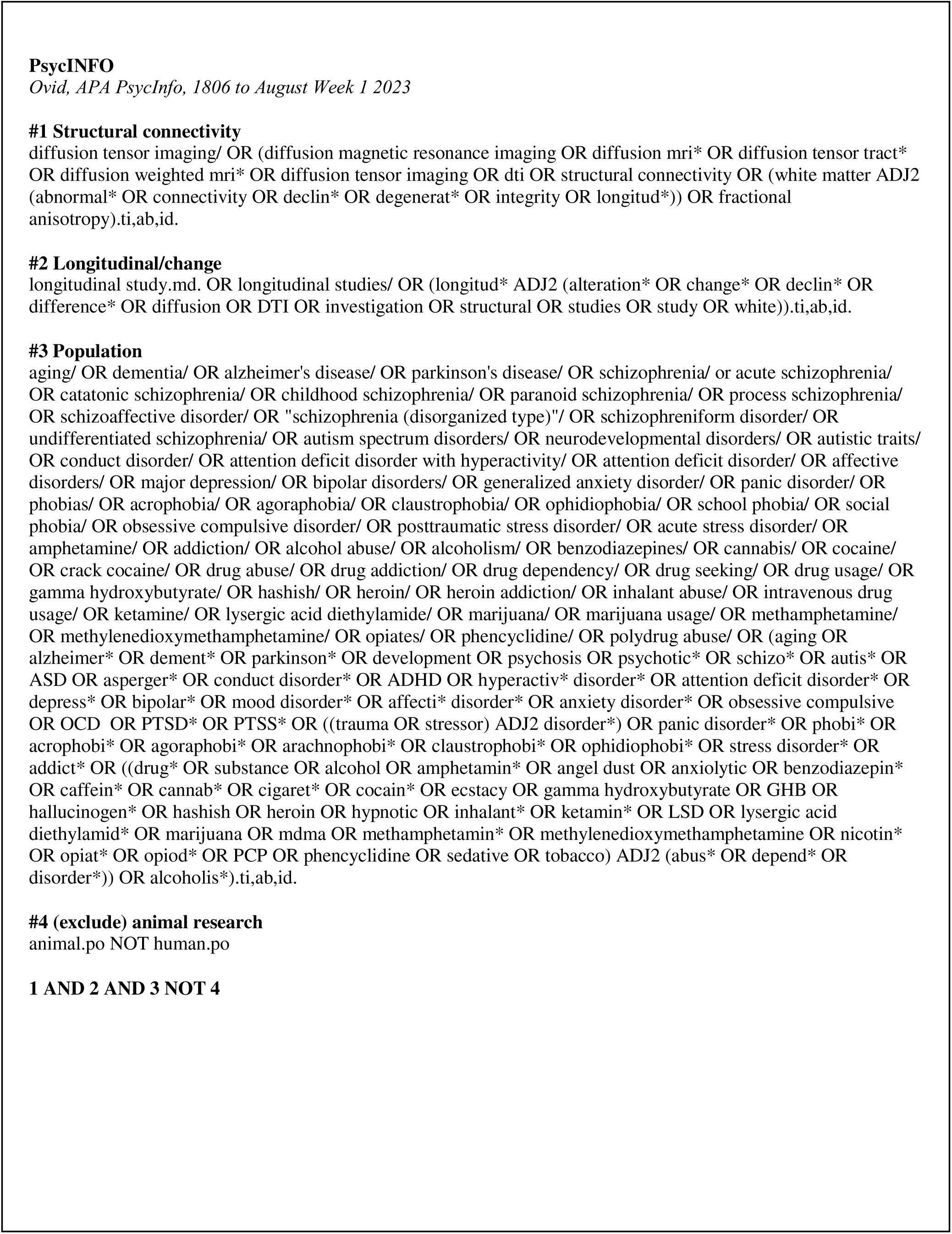
Search syntax from PsycINFO database, last accessed on August 9, 2023.

**Table 6.**
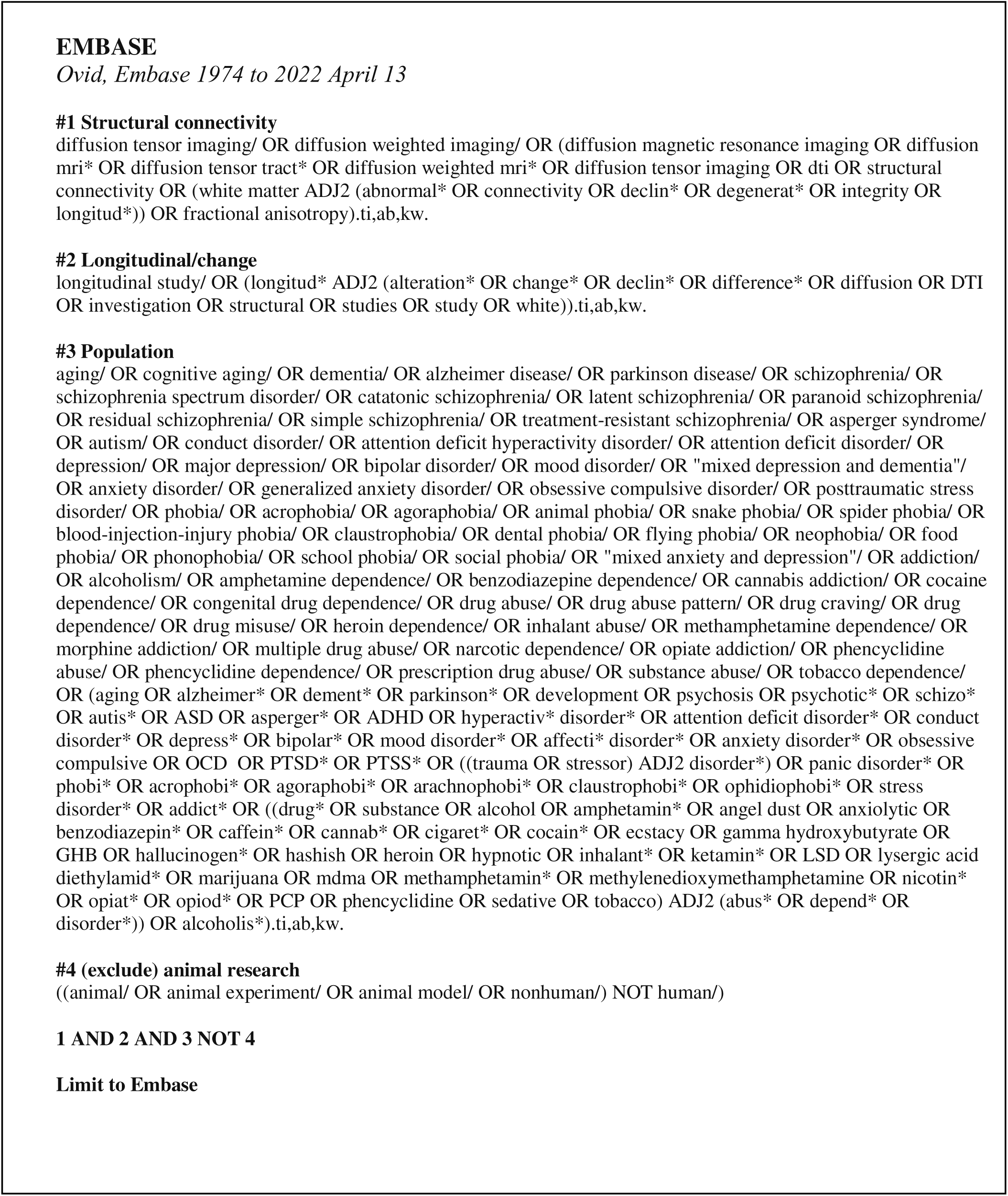
Search syntax from Embase, last accessed on August 9, 2023.

