## Supplementary File for "White matter microstructure changes across the lifespan: a meta-analysis of longitudinal diffusion MRI studies"

***Supplementary 1.1*** *Study Specific Details*

Across several studies the same samples were either totally or partially overlapping. A decision was made to include the studies with the highest number of participants, and so in total 7 studies were excluded to prevent bias. All 29 studies included both male and female subjects and the number of participants per study ranged from 26 (Shakeel et al., 2020) to 488 (Alloza et al., 2018); note that only studies with a minimum of 25 participants per group were included. The minimal interval between baseline and follow-up scan included in the current review was 6 months (Kristensen et al., 2021) and the maximum interval was 5 years (Chiang et al., 2023); note that only studies with a minimum interval of 6 months were included. Twenty five studies measured FA change in healthy individuals, several of these also included patients groups: diabetes (Fox et al., 2018), traumatic brain injury (Graham et al., 2020), ADHD (Chiang et al., 2023) and attenuated syndromes or familial risk of attenuated syndromes (Shakeel et al., 2020). Additionally, one study included patients with HIV (Cole et al., 2018), one study included individuals with alcohol dependency (Pfefferbaum et al., 2014), and one study included individuals who were born prematurely (Adrian et al., 2023). One study included patients with the behavioural variant of frontotemporal dementia (BvFTD), the semantic variant of progressive aphasia (SvPPA) and the non-fluent variant of PPA (NfvPPA) (Staffaroni et al., 2019).

***Supplementary 1.2*** Approximating standard deviation

To approximate the standard deviation (SD) for the remaining studies, Z-scores were first calculated using estimated or reported p-values with the qnorm function in R studio version 1.3.1073 (RStudio Team (2020; <http://www.rstudio.com>). If a two-tailed test was used, p-values were halved before this step. Subsequently, the SD was calculated by dividing the mean FA change by the corresponding Z-score.

***Supplementary 1.3***  Exclusion of three outlier studies

To further assess the robustness of the lifespan trajectory, we conducted an additional sensitivity analysis excluding three studies that appeared to contribute to outlier values (Shakeel et al., 2020; Kruggel et al., 2017; Wolff et al., 2012). These studies provided extreme estimates relative to the rest of the sample. Specifically, the study by Wolff and colleagues (mean age baseline = 0.5 years, ΔFA/year = +0.05) and Shakeel et al., 2020 (mean age baseline = 18 years, ΔFA/year = +0.02) caused the developmental curve to shift upward and altered the estimated age at which the trajectory transitioned from increasing to decreasing FA, as well as changed the rate of decline up until the upper age limit of the sample. The study by Kruggel et al., 2017 (Mean age baseline = 74 years, ΔFA/year = -0.025) also led to a steeper decline in FA towards the upper age limit of the sample (see **Supplementary Figure S1**). Based on these effects, we excluded these three studies from the main model. This decision does not imply that the estimates from these studies are inaccurate, rather, there is currently insufficient data to reliably model their influence within the overall trajectory.

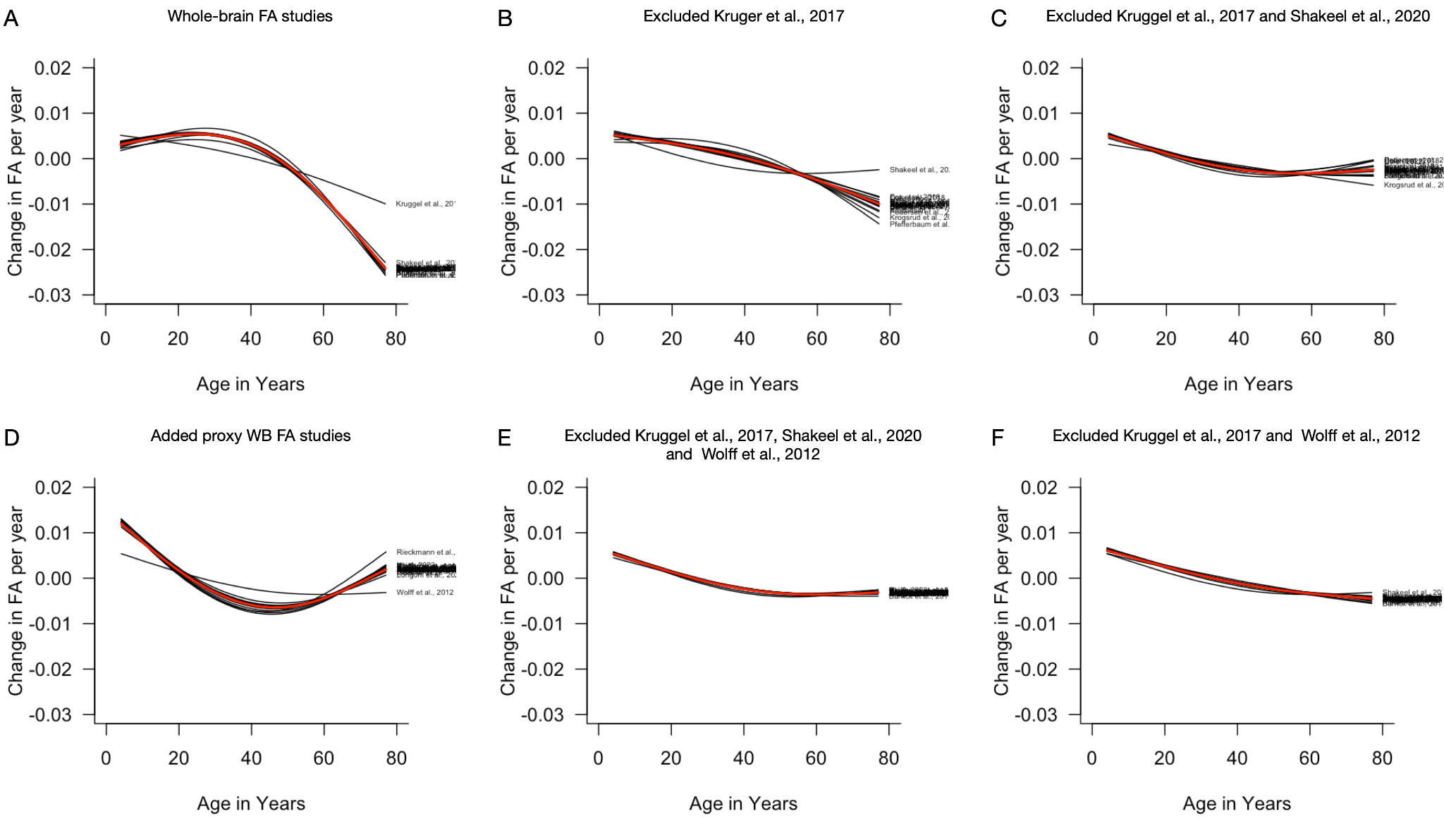

**Figure S1. Sensitivity analysis of annual whole-brain FA change across the lifespan.** Each panel shows the fitted thin plate spline model (3 knots) of age-related change in whole-brain FA based on a meta-regression after excluding one of the studies from the analysis as marked by the label. (**A**) Included all whole-brain FA studies and indicated a strong effect of the study from Kruggel et al., 2017 on the trajectory of fitted curve. (**B**) Excluded the study from Kruggel et al., 2017 (Z-score for change in FA/year of -3.45) from analysis, revealing Shakeel et al., 2020 as a second potential outlier excluding. (**C**) Excluded the studies from Kruggel et al., 2017 and Shakeel et al., 2020, with no further indication of potential outliers. (**D**) After adding studies with proxy for whole-brain FA based on pooled average of individual tract FA, the study from Wolff et al., 2012 was identified as potential outlier (Z-score for change in FA/year of +7.32). (**E**) Excluded the three studies from Kruggel et al., 2017, Wolff et al., 2012 and Shakeel et al., 2020 from analysis, stabilizing the trajectory of the fitted curves in the leave-one-out sensitivity analysis. (**F**) After reintroducing studies back in the analysis it was discovered that the study from Shakeel et al., 2020 had significantly less impact on the trajectory of the curve in the combined whole-brain FA and proxy whole-brain FA analysis. However, because the study was deemed an outlier in the whole-brain FA only studies and the stated change in FA per year was considered unusually high for adolescent development (Z-score of +2.50), and because excluding the study did not have any significant impact on the trajectory of the curve, it was decided to exclude the Shakeel et al., 2020 study from analysis. The analysis in the main text is based on the studies included in Panel C.

***Supplementary 1.4*** Results of a mixed-effect meta-regression model using restricted maximum likelihood (REML) estimation to examine the association between age and annual change in FA.

| **Table S1.** AIC values provided when fitting the data with a thin-plate spline model (with 3, 4 or 5 knots). | | | |
| --- | --- | --- | --- |
| Studies included | 3 knots | 4 knots | 5 knots |
| Category 1 | **-35.32** | -22.33 | -19.67 |
| Category 1 + 2 | **-47.67** | -36.38 | -34.20 |
| Category 1 + 2 + 3 | **-91.02** | -81.32 | -71.45 |
| Category 1 + 2 + 3 + 4 | **-100.88** | -96.61 | -86.88 |
| Category 1 + 2 + 3 + 4 + 5 | **-176.32** | -166.62 | -158.12 |

*Note*: Addition of category 5 are the studies that measured FA change across individual tracts. A pooled estimate of annual FA change across all reported tracts was calculated.

With all categories included the addition of the thin-plate spline terms modelling age-dependency on annual FA change significantly improved model fit (QM(2) = 92.13, *p* < .001), indicating a non-linear relationship between age and annual FA change. Both spline terms were highly significant (*p* = .016 and *p* < .001), supporting a robust non-linear association. Across childhood and adolescence, FA increased, with the rate of increase slowing into early adulthood. Between approximately ages 20 and 35, changes in FA were not statistically significant. This was followed by a significant decline in FA between ages 36 and 50. The decreases plateaued by the age of 51 up until the upper end of the age range (77 years).

**Table S2.** Results of a mixed-effect meta-regression model using restricted maximum likelihood (REML) estimation to examine the association between age and annual change in FA. Table displays the model terms, estimated coefficients, standard errors, z-values, p-values and 95% confidence intervals and statistical outputs.

| **Model term** | **Estimate** | **SE** | **z-value** | **p-value** | **95% CI** |
| --- | --- | --- | --- | --- | --- |
| Intercept | -0.0005 | 0.0004 | -1.3517 | .177 | [-0.0013: 0.0002] |
| Age spline component 1 | -0.0053 | 0.0022 | -2.4048 | .016 | [-.0097: -0.0010] |
| Age spline component 2 | -0.0028 | 0.0005 | -5.8731 | < .001 | [-0.0038: -0.0019] |
| **Statistics** | | | **Value** | | |
| Number of studies | | | 26 | | |
| Log-likelihood | | | 92.16 | | |
| AIC | | | -176.32 | | |
| BIC | | | -171.77 | | |
| tau² (Between-study variance) | | | 2.59e-06 SE(9.41-07) | | |
| tau | | | 0.0016 | | |
| I² (Residual heterogeneity) | | | 99.88% | | |
| R² (Heterogeneity explained by moderators) | | | *0.00%* | | |
| Test for residual heterogeneity (QE, df = 23) | | | *18,628.54, p < .0001* | | |
| Test of moderators (QM, df = 2) | | | 92.13, *p* < .001 | | |

***Supplementary 1.5*** Categorisation of studies based on available data

A sensitivity analysis was conducted to assess the robustness of results from studies with approximated parameters. Studies were classified into four categories based on the availability and completeness of their reported statistical information. Category 1 included studies that reported both a point estimate and either a SD or a p-value. Category 2 comprised studies where the point estimate was approximated (e.g. from a figure), but the p-value was reported and used to estimate the SD. In Category 3, studies provided a point estimate, but the SD had to be estimated using a conservative p-value. Finally, Category 4 included studies for which both the point estimate and the SD or p-value had to be approximated. Each category was added in succession to test whether the trajectory of FA change varied (See **Supplementary Figure S2**).

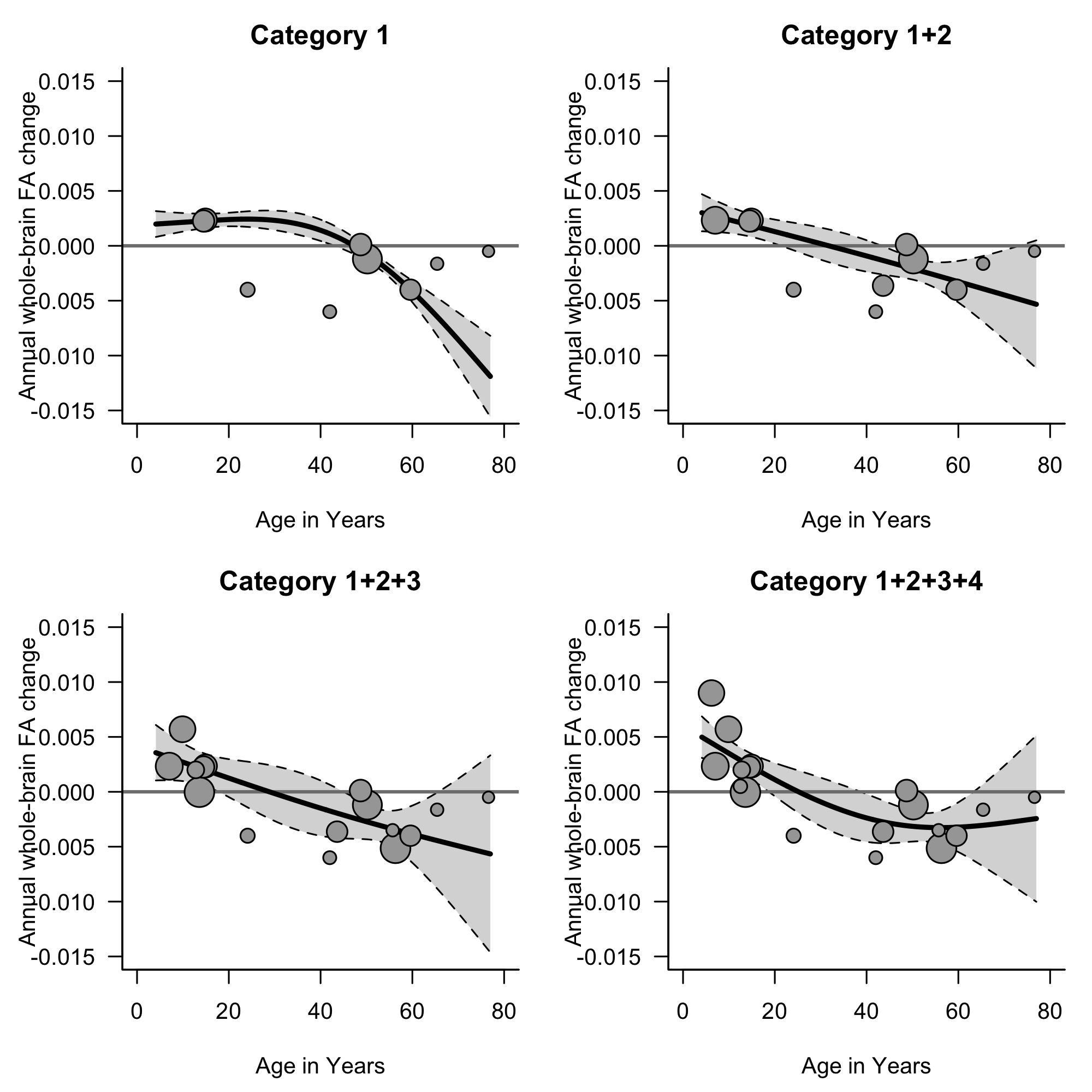

**Figure S2.** **Meta-analysis of whole-brain FA change across the lifespan**. Annual FA change per year represents the FA at timepoint 2 minus FA at timepoint 1, divided by the number of years between timepoints. Each circle represents a cohort, with circle size proportional to cohort size (ranging from N = 32 to N = 213). Data were pooled across groups within each study. Studies were classified into four categories: Category 1 reported both point estimates and SD or p-values (blue circles); Category 2 required approximation of the point estimate but reported p-values to estimate SD (green circles); Category 3 reported point estimates but used a conservative p-value to estimate SD (purple circles); Category 4 required approximation of both the point estimate and SD or p-values (grey circles). The addition of each category is visualised in the plots. A thin-plate spline model was used to fit the data, shown with 95% confidence intervals

***Supplementary 1.6***  Inclusion of studies measuring FA in individual tracts

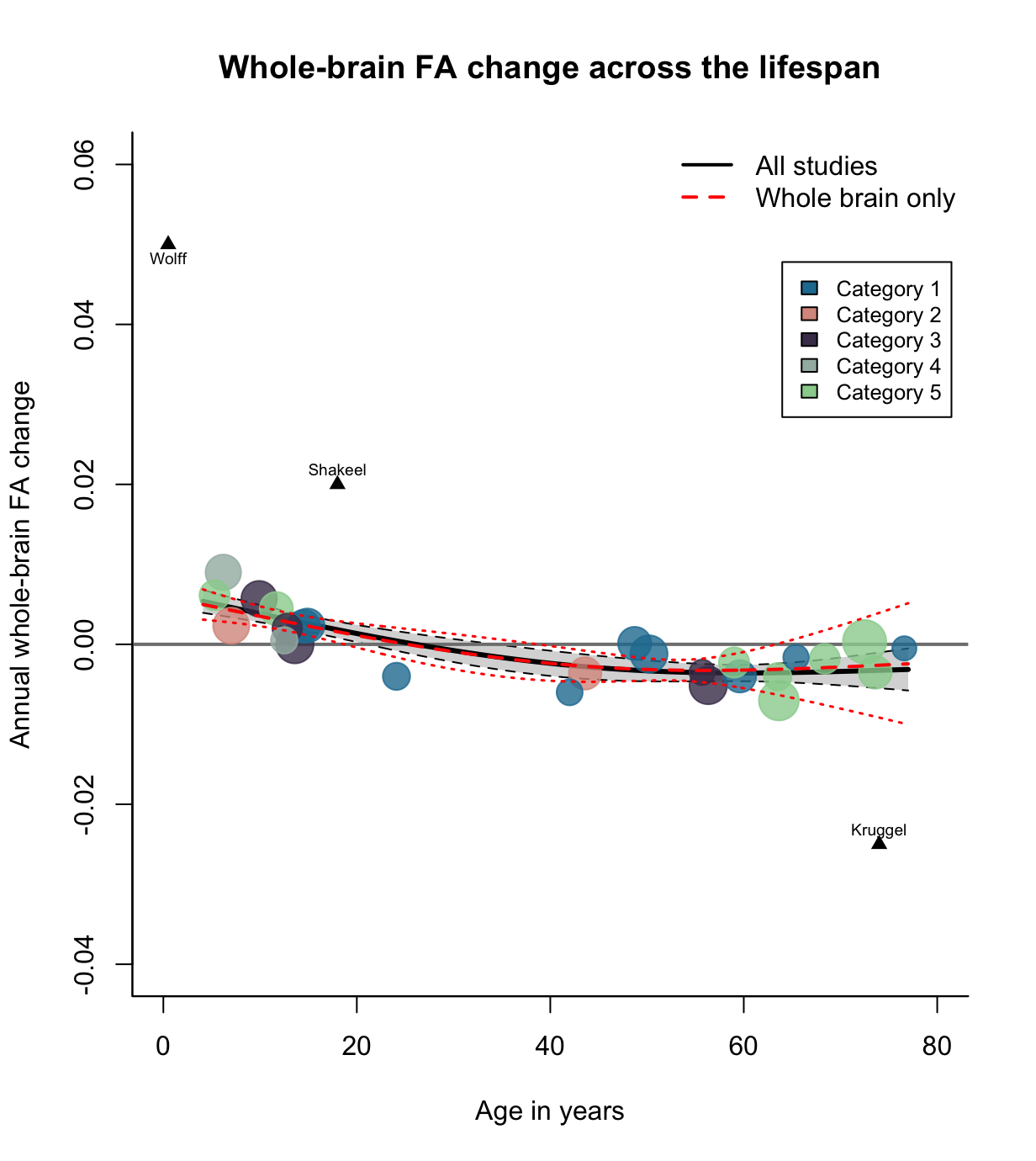

**Figure S3. Meta-analysis of whole-brain FA change across the lifespan, including both studies that measured whole-brain FA directly and those that reported tract-wise values from which whole-brain estimates were approximated.** Three studies (Shakeel et al., 2020, Kruggel et al., 2017 and Wolff et al., 2012) were excluded from the thin plate spline modelling due to extreme values but are shown here for visualisation purposes. Annual FA change per year represents the FA at timepoint 2 minus FA at timepoint 1, divided by the number of years between timepoints. Each circle represents a cohort, with circle size proportional to cohort size (ranging from N = 32 to N = 213). Studies were classified into four categories: Category 1 reported both point estimates and SD or p-values (blue circles); Category 2 required approximation of the point estimate but reported p-values to estimate SD (green circles); Category 3 reported point estimates but used a conservative p-value to estimate SD (purple circles); Category 4 required approximation of both the point estimate and SD or p-values (grey circles); Category 5 are the studies that measured FA change across individual tracts. A pooled estimate of annual FA change across all reported tracts was calculated. A thin-plate spline model was used to fit the data, shown with 95% confidence intervals. Two spline models are shown: one including all studies (solid black line), and one including only studies that directly reported whole-brain FA (dashed red line).

***Supplementary 1.7*** Comparison of whole-brain FA change across the lifespan in healthy controls versus full sample

A similar pattern of annual FA change was found when only investigating healthy control populations. Across childhood, adolescence FA increased, with the rate of increase slowing into early adulthood. Between approximately ages 25 and 36, changes in FA were not statistically significant. This was followed by a decline in FA between ages 37 and 62. The decreases plateaued by ages 65 up until the upper limit of the age range (77 years) (**Supplementary** **Figure S4**).

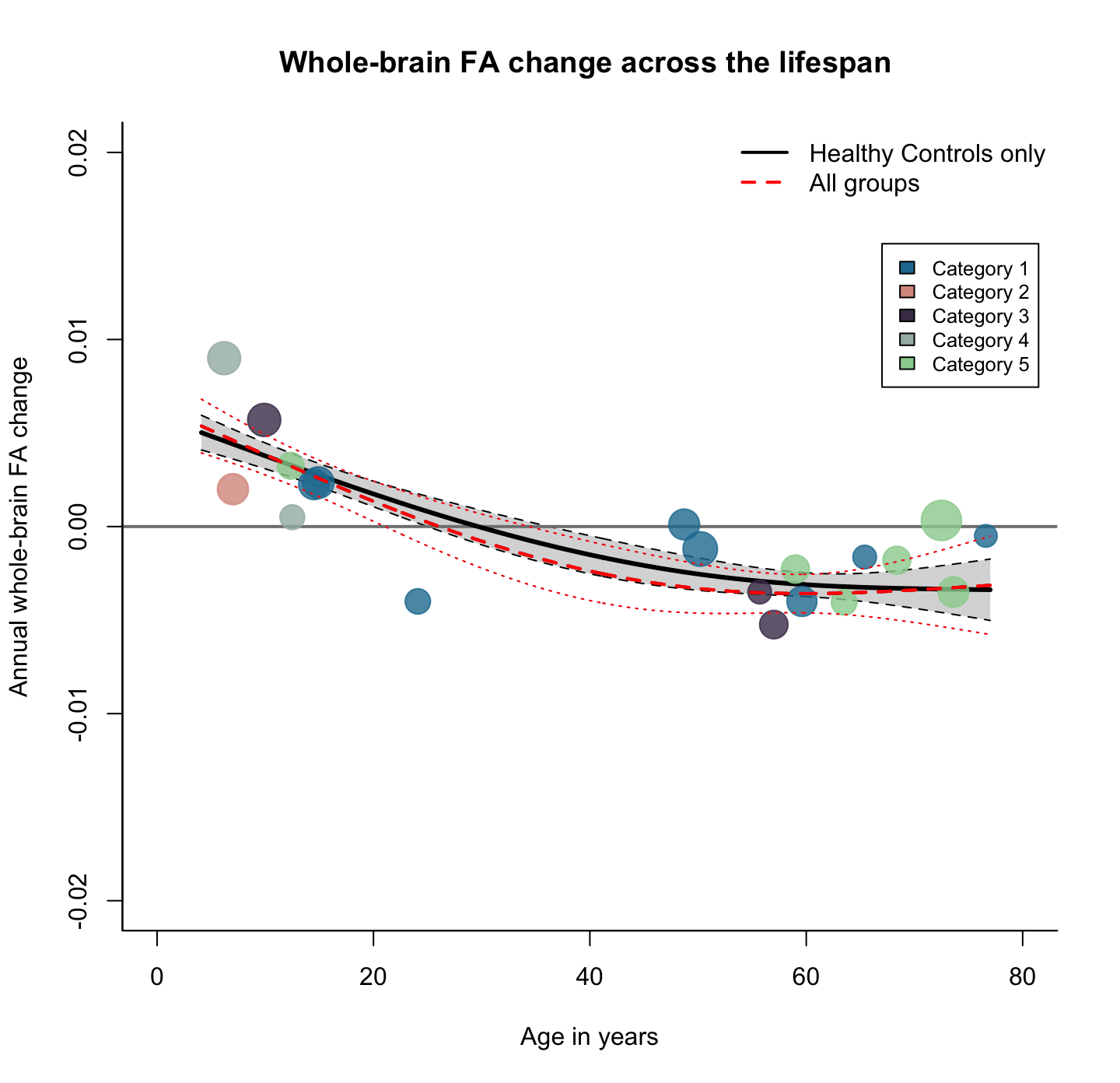

**Figure S4.** **Meta-analysis of whole-brain FA change in healthy populations versus the full sample across the lifespan**. Annual FA change per year represents the FA at timepoint 2 minus FA at timepoint 1, divided by the number of years between timepoints. Each circle represents a cohort, with circle size proportional to cohort size (ranging from N = 32 to N = 203). Studies were classified into four categories: Category 1 reported both point estimates and SD or p-values (blue circles); Category 2 required approximation of the point estimate but reported p-values to estimate SD (green circles); Category 3 reported point estimates but used a conservative p-value to estimate SD (purple circles); Category 4 required approximation of both the point estimate and SD or p-values (grey circles); Category 5 included studies that measured FA change across individual tracts, from which pooled estimate of annual FA change was calculated. A thin-plate spline model was used to fit the data, shown with 95% confidence intervals. Two spline models are plotted: one including only healthy populations (solid black line and data points), and one including the full sample (dashed red line).

| **Table S3.** The list of individual tracts which were included in the averaged annual FA change across tracts per study. | | |
| --- | --- | --- |
| Study | Tracts | N of regions/tracts |
| Vik et al., 2015 | Thalamus-superior frontal, caudate-superior frontal, putamen-superior frontal, caudate-medial orbitofrontal, putamen-medial orbitofrontal, accumbens-medial orbitofrontal. Caudate-lateral orbitofrontal, putamen-lateral orbitofrontal, putamen-par triangularis, thalamus-rostral middle frontal, thalamus-rostral middle frontal, caudate-rostral middle frontal, putamen-rostral middle frontal (for both left and right hemispheres) | 19 |
| Barrick et al., 2010 | Internal capsule- posterior limb, internal capsule anterior limb, cingulum-anterior, cingulum-posterior, external capsulse, superior corona radiation, (for both left and right hemispheres), genu, body and splenium of corpus callosum. | 15 |
| Staffaroni et al., 2019 | Frontal (superior longitudinal fasciculus, cingulum bundle), temporal (fornix, inferior longitudinal fasciculus, sagittal stratum, hippocampal portion of the cingulum), uncinate fasciculus (for both left and right hemispheres), and genu of corpus callosum,  right and left frontal (superior longitudinal fasciculus, cingulum bundle) and temporal (fornix, inferior longitudinal fasciculus, sagittal stratum, hippocampal portion of the cingulum) lobes, along with the uncinate fasciculus and genu of the corpus callosum | 15 |
| Alloza et al., 2018 | Arcuate fasciculus, thalamic radiation, cingulum, uncinate fasciculus, inferior longitudinal fasciculus, genu and splenium of corpus callosum | 7 |
| Adrian et al., 2023 | Corpus callosum, thalamic radiation, cingulum, corticospinal, inferior fronto-occipital fasciculus, inferior longitudinal fasciculus, superior longitudinal fasciculus (parietal and temporal) and uncinate fasciculus. | 9 |
| Rieckmann et al., 2016 | Inferior fronto-occipital fasciculus, superior frontal occipital fasciculus, corona radiata (left and right hemispheres), internal capsule, cingulum, cingulum hippocampus, corona radiata (anterior), body, splenium and genu of corpus callosum | 12 |
| Chiang et al., 2023 | Corpus callosum (prefrontal cortex, sensorimotor cortex, parietal cortex, temporal cortex) Arcuate fasciculus, superior longitudinal fasciculus (I, II, III), frontal aslant, perpendiculus fasciculus, cingulum (main body, hippocampus), stria terminalis, uncinate fasciculus, inferior fronto-occipital fasciculus, inferior longitudinal fasciculus, fronto-striatal (prefrontal cortex, precentral gyrus) thalamic radiation (prefrontal cortex, sensorimotor cortex, auditory radiation, optic radiation), corticospinal, (left and right hemispheres), fornix and splenium of corpus callosum. | 44 |
| Coelho et al., 2021 | Two clusters. Cluster 1 (body of corpus callosum and right superior and posterior corona radiata). Cluster 2 (genu, body, splenium of corpus callosum, anterior limb of internal capsule, anterior, superior and posterior corona radiata, external capsule, and superior longitudinal fasciculus) | 12 |

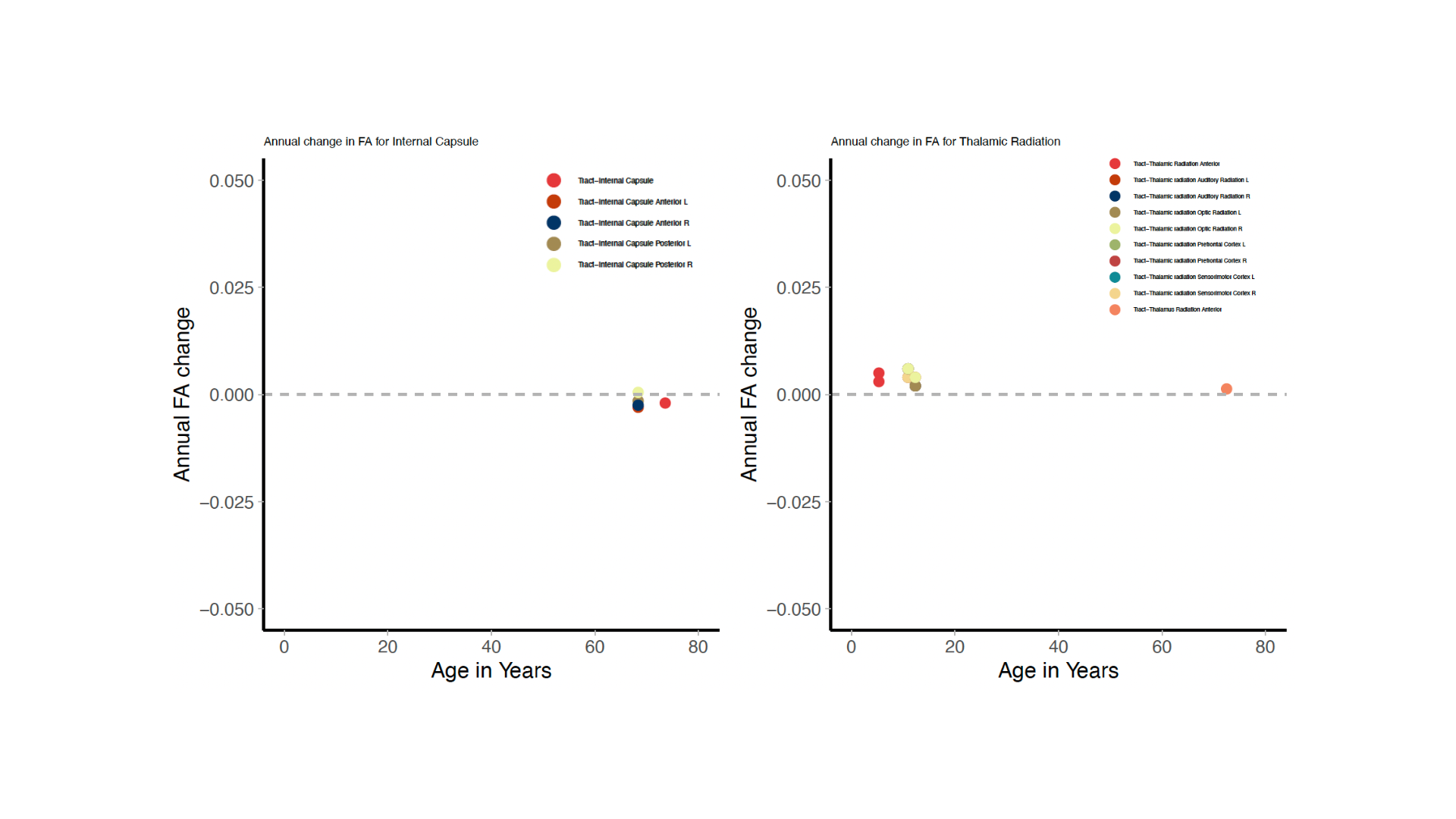

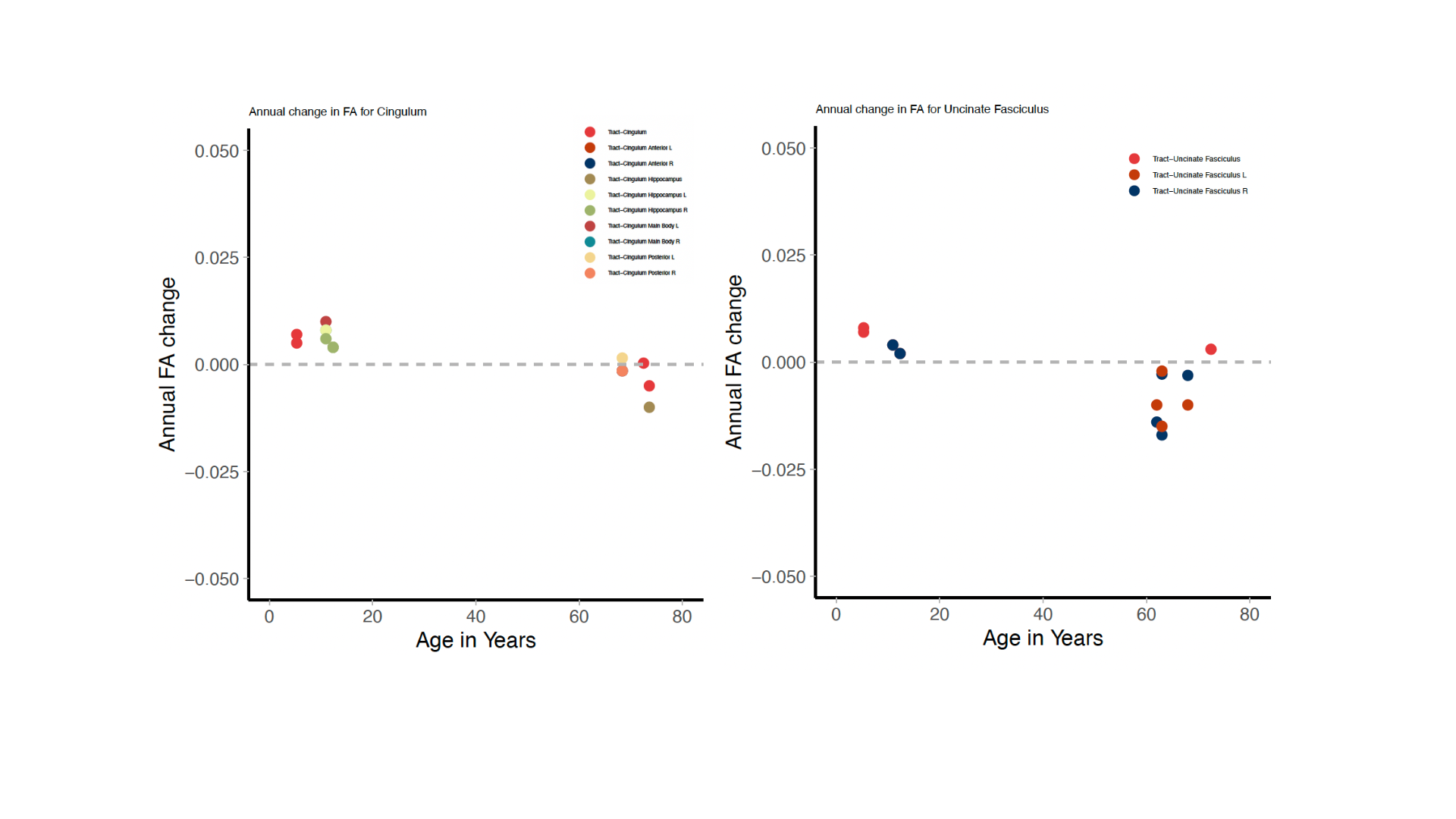
**Figure S5.** Plots per individual tract.

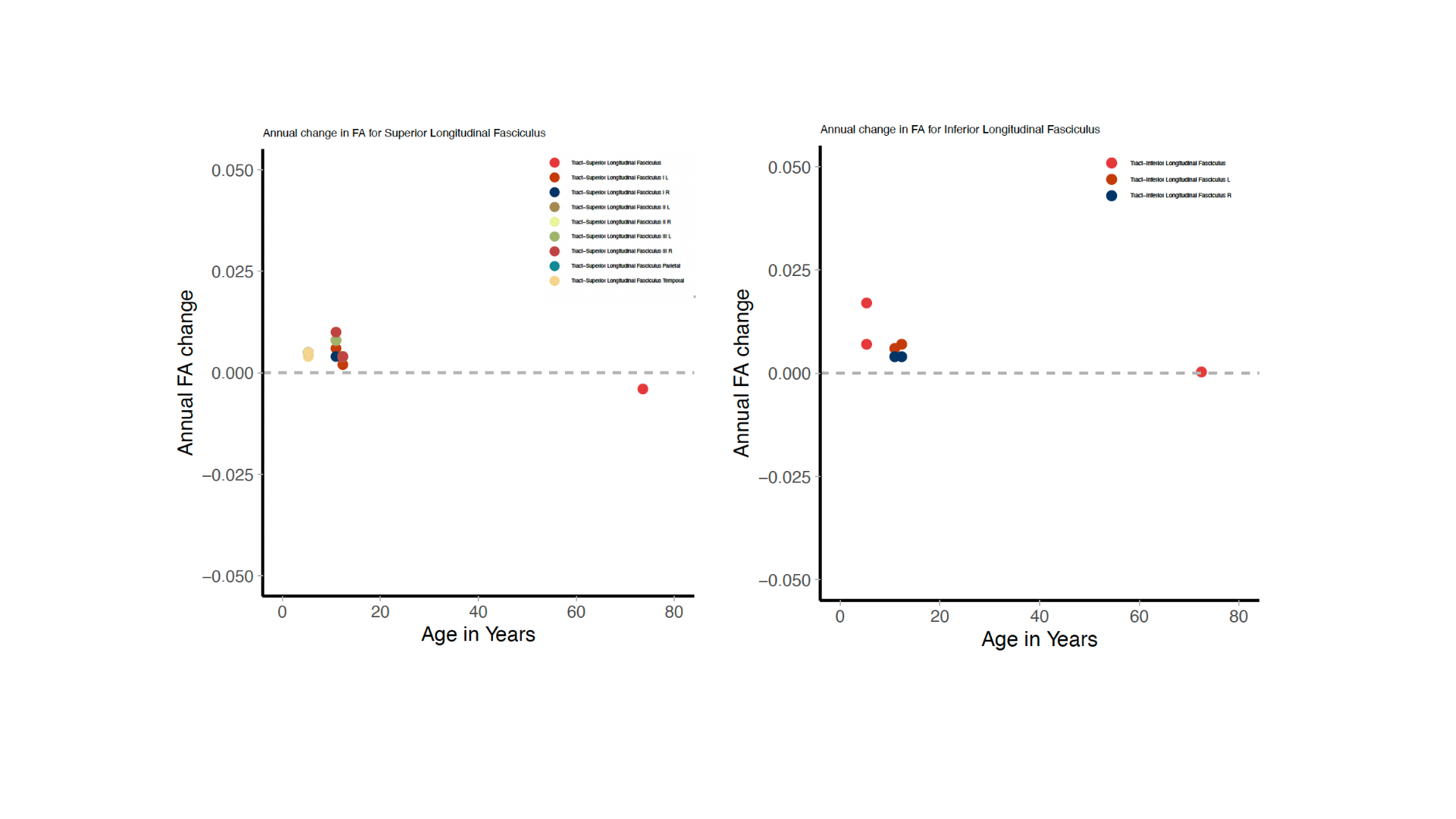
